# Psi RNA-specific Binding Promotes HIV-1 Gag Conformational Change Critical for Immature Viral Particle Assembly

**DOI:** 10.64898/2026.07.18.739361

**Authors:** Yehong Qiu, Puja Banerjee, Kaylee Grabarkewitz, Vicki H. Wysocki, Ioulia Rouzina, Gregory A. Voth, Karin Musier-Forsyth

**Author notes:** Karin Musier-Forsyth (Corresponding author). equal contributions (Yehong Qiu, Puja Banerjee).

## Abstract

The immature HIV-1 virion is assembled by the Gag polyprotein using inositol hexakisphosphate (IP6) as an essential assembly co-factor. Gag binds the genomic RNA Psi packaging signal via the nucleocapsid (NC) domain and associates with the plasma membrane via the matrix (MA) domain. Previous studies revealed that Gag exists in both compact (C) and extended (E) conformational states in solution. Only E-Gag formed virus-like particles with the correct size and IP6 shifted the equilibrium of DNA-bound Gag to the E state. The influence of specific RNA elements on this conformational change is unknown. In this work, a dual dye-labeled Gag was prepared for probing the effect of RNA binding on Gag conformation using Förster resonance energy transfer (FRET). In low salt and in the absence of other factors, Gag was primarily in the C state. Psi RNA binding induced a more significant FRET decrease than binding to non-Psi RNAs, consistent with a shift to E-Gag. IP6 alone also promoted the E-Gag state in the absence and presence of RNA. Atomistic molecular dynamics simulations are consistent with and provide detail into the role of NC-Psi RNA binding in the conformational switch of C-Gag to assembly-competent E-Gag. Simulations also showed that this switch is driven by capsid (CA) linker domain orientational flexibility and MA-CA unbinding dynamics. Thus, the highly flexible multi-domain Gag polyprotein leverages both viral and host cell factors to sample and stabilize distinct conformations, thereby orchestrating the viral assembly process.

## Introduction

Human immunodeficiency virus type 1 (HIV-1) is the causative agent of acquired immunodeficiency syndrome (AIDS). Effective antiretroviral therapies are available; however, HIV-1 can readily acquire drug resistance due to its high mutation rate and no cure exists. Therefore, therapeutics that target novel steps in the viral lifecycle such as virion assembly are urgently needed (Nastri et al. 2023).

In the viral assembly process, the dimeric HIV-1 genomic RNA (gRNA), along with viral proteins and cellular factors are packaged into an immature virion formed by approximately 2500 Gag molecules, which co-assemble with the Gag-Pol polyprotein (Shehu-Xhilaga et al. 2001; Chen et al. 2009). Although all components are essential for maximum infectivity, packaging of two copies of gRNA is especially important (Moore and Hu 2009). One critical function of Gag is its role in selectively packaging the dimeric gRNA in the presence of a vast excess of cellular RNAs in the cytoplasm (Comas-Garcia et al. 2016). This occurs, in part, due to the interaction between the nucleocapsid (NC) domain of Gag and the Psi packaging signal located in the 5‘UTR. Although Gag demonstrates specific binding to HIV-1 gRNA relative to other RNAs in cells, under physiological conditions *in vitro*, Gag’s binding affinity to Psi and non-Psi RNAs is similar (Webb et al. 2013; Comas-Garcia et al. 2017). Thus, in addition to binding affinity, other factors such as cellular localization (Maldonado et al. 2020; Lambert et al. 2024), number of 5‘ guanosine residues (Masuda et al. 2015; Kharytonchyk et al. 2016; Nikolaitchik et al. 2021) and Gag conformation have been proposed to play a role in specific gRNA selection (Olson and Musier-Forsyth 2019).

The domains of the Gag polyprotein include matrix (MA), capsid (CA), spacer peptide 1 (SP1), NC, spacer peptide 2 (SP2) and p6 all connected by flexible linkers. MA is myristoylated on Gly2, which facilitates Gag targeting to the inner leaflet of the host cell plasma membrane (PM). Membrane binding occurs via interactions mediated by both the myristoyl moiety and the MA highly-basic region (HBR) with phosphatidylinositol-(4,5)-bisphosphate (PIP2)-rich regions of the PM (Saad et al. 2006). The HBR also binds to RNAs, including cellular tRNAs that are involved in the regulation of PM binding (Thornhill et al. 2020). CA has two structured domains connected by a linker. The C-terminal domain (CTD) residues W316 and M317 are key for mediating Gag-Gag interactions (von Schwedler et al. 2003). Upon Gag oligomerization, the entire SP1 region together with the last eight residues of the CA_CTD_ form a 6-helix bundle (6HB) to stabilize the Gag hexamer (Datta et al. 2016). Another important factor for Gag assembly is inositol hexakisphosphate (IP6), which is present in mammalian cells at concentrations between 10 µM and 1 mM (Shamsuddin 1999). In the immature Gag lattice, a single IP6 stabilizes two rings in the 6HB by binding to K290 and K359 (Dick et al. 2018). In the mature CA lattice, two IP6 molecules bind to R18 and K25 to stabilize two different rings in the CA_CTD_ (Yu et al. 2020; Kleinpeter et al. 2024). NC is a highly basic protein that encodes two zinc finger domains responsible for gRNA binding and chaperone function (Levin et al. 2010; Mouhand et al. 2020).

The Gag polyprotein has been reported to have multiple conformations in solution. An early small-angle neutron scattering study of a monomeric Gag variant with W316A/M317A mutations (WM-Gag) demonstrated that Gag does not form a single conformation in solution, consistent with its high flexibility (Datta et al. 2007a). In the immature particle, Gag molecules align in parallel orientation in an elongated, rod-shaped manner. Thus, Gag must undergo a significant conformational change during assembly. A subsequent study by Rein and co-workers concluded that both nucleic acids and lipids are required for Gag to adopt an elongated conformation (Datta et al. 2011). A single-molecule Förster resonance energy transfer (smFRET) study of Gag concluded that two major states of Gag, extended (E) and compact (C), exist in solution (Munro et al. 2014). Only E-Gag formed viral-like particles (VLPs) comparable in size similar to immature virions in cells. This study showed that the addition of a polyA_25_ DNA oligonucleotide to stoichiometric amounts of Gag led to formation of small VLPs formed by C-Gag molecules, while in the presence of both DNA and IP6, larger VLPs formed by E-Gag were observed. A computational study predicted that Gag alone exists in two alternative conformations, with and without intra-Gag contacts, respectively (Su et al. 2018). A separate study using molecular dynamics (MD) simulations predicted that C-Gag is stabilized by specific interactions between residues in the CA_CTD_ and MA (Lin et al. 2019). Using fluorescence imaging, a fraction of Gag molecules was shown to adopt the C state in cells (Zeiger et al. 2024). In this work, various predictions of the computational models were tested and these studies were consistent with C-Gag stabilization by CA-MA interactions. Mutants that reduced C-Gag formation also reduced particle production and infectivity, suggesting a role for C-Gag in the viral lifecycle.

Using native mass spectrometry, Psi RNA has been shown to promote the dimerization of WT Gag but not WM-Gag (Sarni et al. 2020); however, whether specific RNA binding alone induces or facilitates a conformational change from C-Gag to E-Gag is unknown. The results of fluorescence anisotropy salt-titration binding assays indirectly supported a role for specific RNA binding in Gag conformation. These assays measured Z_eff_, the number of cations displaced from the RNA upon protein binding, reflecting the electrostatic interaction (Rye-McCurdy et al. 2015). The Z_eff_ measured for Gag binding to Psi RNA was ∼5, a value similar to that of the NC domain binding to either Psi or non-Psi RNA (Webb et al. 2013). This supports an NC-only binding mode for Gag binding to Psi RNA. In contrast, the high Z_eff_ (∼9) measured for Gag binding to non-Psi RNA suggested that in this case, the Gag-RNA interaction occurred with both MA and NC, likely in the C state.

In this study, our goal was to directly probe the conformation of full-length monomeric Gag upon binding to a variety of RNAs. Gag was dually labeled at the N-terminus with Cy3 and near the C-terminus with Cy5 to enable ensemble FRET measurements. The FRET efficiency value in the absence of RNA suggested that Gag is primarily in a C- conformation in solution. A decrease in FRET was measured upon binding of all RNAs tested, with Psi RNA inducing the maximum decrease, implying the strongest shift toward the E state. The Gag assembly co-factor IP6 also shifted the equilibrium from the C- to E-Gag state, even in the absence of RNA. All-atom MD simulations were consistent with the experimental results and further revealed a molecular-level mechanism of C-Gag to E-Gag transition in the presence of specific RNA binding to the NC domain of Gag. Mass photometry (MP) studies showed a significantly higher population of Gag multimers bound to Psi RNA than to a non-Psi RNA under identical solution conditions. Taken together, the experimental and *in silico* results highlight the critical role of specific Psi RNA binding in Gag conformation and oligomerization, with important implications for retroviral assembly.

## Results

### Double-labeling of Gag and quantitative analysis of the dual-labeled Gag FRET signal

The scheme developed to label Gag with a Cy3 (donor) and Cy5 (acceptor) is shown in Figure 1A. As described in the Methods, Sortase was used to ligate the Cy3-conjugated LEPTGG peptide to Gag containing an N-terminal triglycine motif. A Lys-Cys-Lys motif was engineered near the Gag C-terminus ^494^KCK^496^ to provide a highly-reactive C495 for Cy5 maleimide conjugation (Hermanson 1996). We anticipated that Gag alone would exhibit significant FRET, consistent with C-Gag, and that a conformational shift toward E-Gag would decrease the FRET signal (Figure 1B). Following the optimized protocol described in the Methods, a final labeling efficiency of ∼55% was consistently obtained for each dye and denaturing PAGE confirmed the purity and labeling of the samples (Figure S1A).

**Figure 1.**
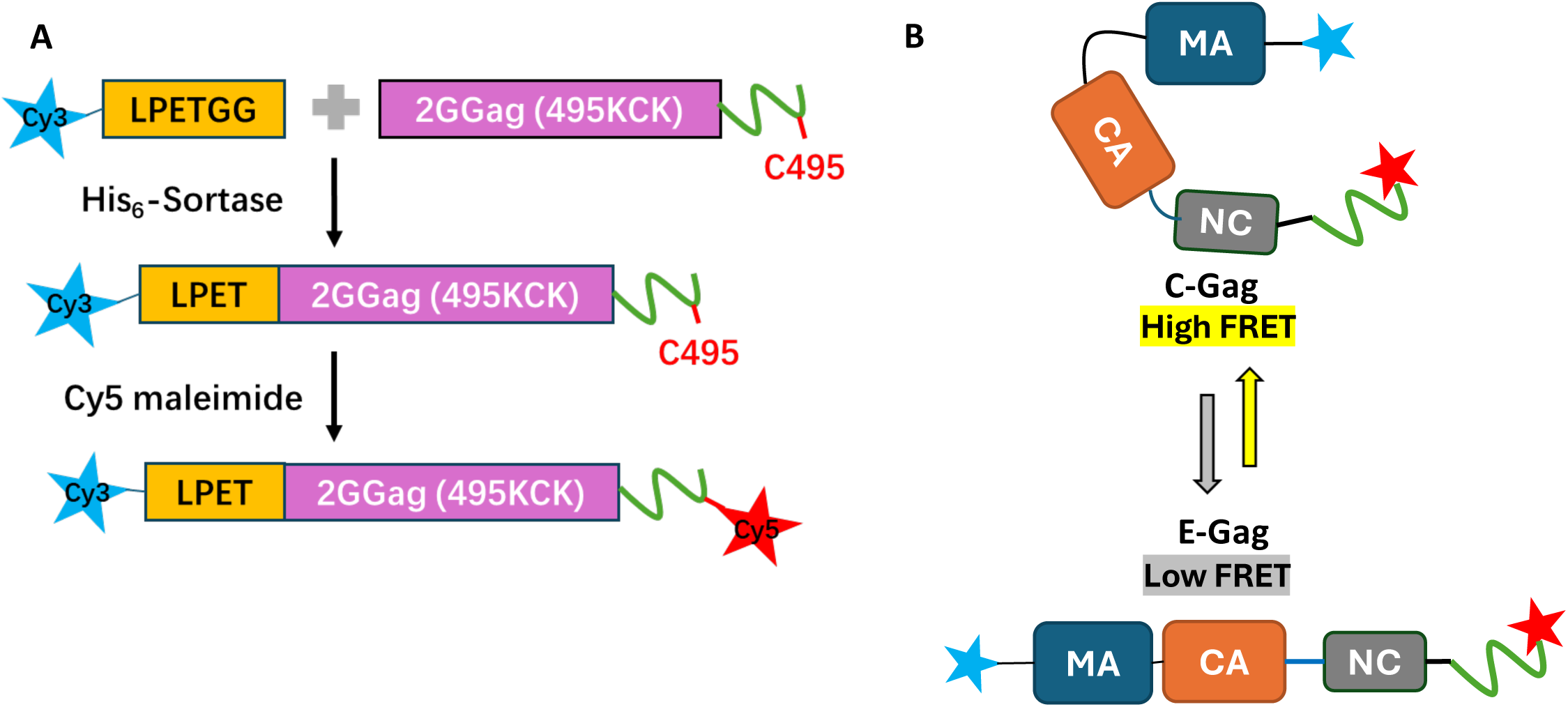
Gag labeling scheme and major conformational states. A) Workflow of Gag dual labeling with Cy3 and Cy5 fluorophores. Two additional glycine residues were inserted after the start codon of Gag to generate an N-terminal triglycine motif required for Sortase-mediated ligation. The cysteine at 495 was activated by imbedding it within a KCK motif. B) Schematic of the two conformation states expected for WT and WM-Gag: compact (C-Gag) and extended (E-Gag).

Published single-molecule FRET results show that Gag exists in either C- or E- states with FRET values of 1 or 0, respectively (Munro et al. 2014). In low NaCl, the maximum FRET efficiency (FE_max_) value (calculated using the *(ratio)*_A_ method, as described in the Methods (eq. 1)) we observe for dual-labeled Gag is ∼0.55 (Figure 2A). This value is consistent with the majority (∼95%) of labeled molecules being doubly-labeled and in the C state. Normalizing our observed FE values by this FE_max_ allows for an estimate of the fraction of Gag molecules in the C-state, *θ_C_*, according to (eq. 2), and of the free energy of the C-to-E transition *g_CE_* (eq. 3 and eq. 4).

**Figure 2.**
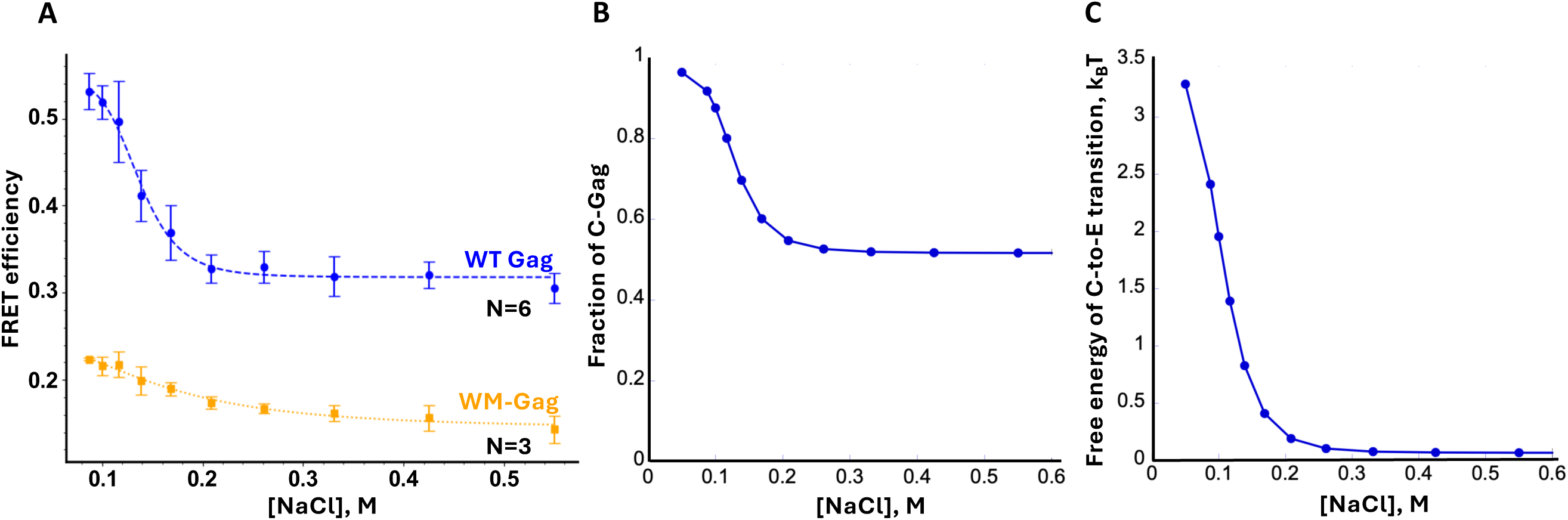
FRET efficiency as a function of NaCl concentration for dual-labeled Gag. A) FRET efficiency of 20 nM WT Gag (blue) and WM Gag (orange) as a function of [NaCl]. B) and C) are the fraction of WT C-Gag and the free energy of the C-to-E transition vs [NaCl] calculated from FRET efficiencies according to Eq. 2 and 4, respectively. Curves are drawn to guide the eye and are not fitted to the data.

### Higher salt stabilizes the E-Gag conformation

We next measured the FE of Gag as a function of increasing NaCl concentration. The FE decreased with increasing NaCl concentration from the maximum value of ∼0.55 at ∼100 mM NaCl to a plateau value of ∼0.32 at [NaCl] ≥ 200 mM. The fraction of C-Gag (calculated according to eq. 2) decreased from 0.95 at 100 mM NaCl to 0.58 at 200 mM NaCl (Figure 2B), and the corresponding transition free energy *g_CE_* (calculated according to eq. 4) decreased from 3.0 *k_B_T* to 0.1 *k_B_T* (Figure 2C). The fact that 200 mM NaCl is sufficient to disrupt the intra-Gag interactions indicates that C-Gag is stabilized by a relatively weak electrostatic component. We also measured FE values of a monomeric WM-Gag variant wherein two hydrophobic residues in the CA C-terminal domain of Gag are mutated to alanine (W184A/M185A). Despite a very similar labeling efficiency, the FE_max_ value for WM-Gag was ∼ 0.22 at the lowest salt concentrations tested, consistent with a primarily E-state conformation (Figure 2A). This result implies that the WM site participates in the intra-Gag interaction that stabilizes the C-Gag conformation, in agreement with observations in cells (Zeiger et al. 2024). At high salt, approximately half of the wild-type (WT) Gag molecules remain in the C-state and the C-to-E transition free energy is close to zero, *g_CE_*∼0 (see Figures 2B and 2C). This implies that the free energy of intra-Gag contacts stabilizing the C-state is equal to the difference in the entropic free energy of the E-state (higher entropy) relative to the C-state.

### Gag binding to Psi RNA promotes a greater conformational shift towards E-Gag than non-Psi RNAs

Based on the salt-sensitivity of the observed FRET, we carried out the following studies at 100 mM NaCl to mimic physiological conditions and to maximize the window for observing a potential FRET change upon factor binding. Psi RNA and two non-Psi RNAs were tested for their effect on the conformation of dual-labeled Gag. The non-Psi RNAs chosen included another element from the HIV-1 gRNA 5‘UTR known as the transactivation response element-polyadenylation hairpin (TARpA) and human tRNA^Lys3^, the primer for HIV-1 reverse transcription (Figure S2). The results of titrating these RNAs into 20 nM Gag in 100 mM NaCl are shown in Figure 3A. Plots of the FE as a function of RNA concentration showed that all RNAs induced a decrease in FRET, with Psi displaying the largest change (Figure 3A). The corresponding changes in the fraction of C-Gag are plotted in Figure 3B. These data support our hypothesis that Psi RNA promotes the conformational change of Gag from the C state to the E state in the absence of any other factors, with non-Psi RNAs having a reduced effect. The impact of RNA binding on inducing the C-to-E conformational transition in Gag followed the order Psi > tRNA^Lys3^ > TARpA.

**Figure 3.**
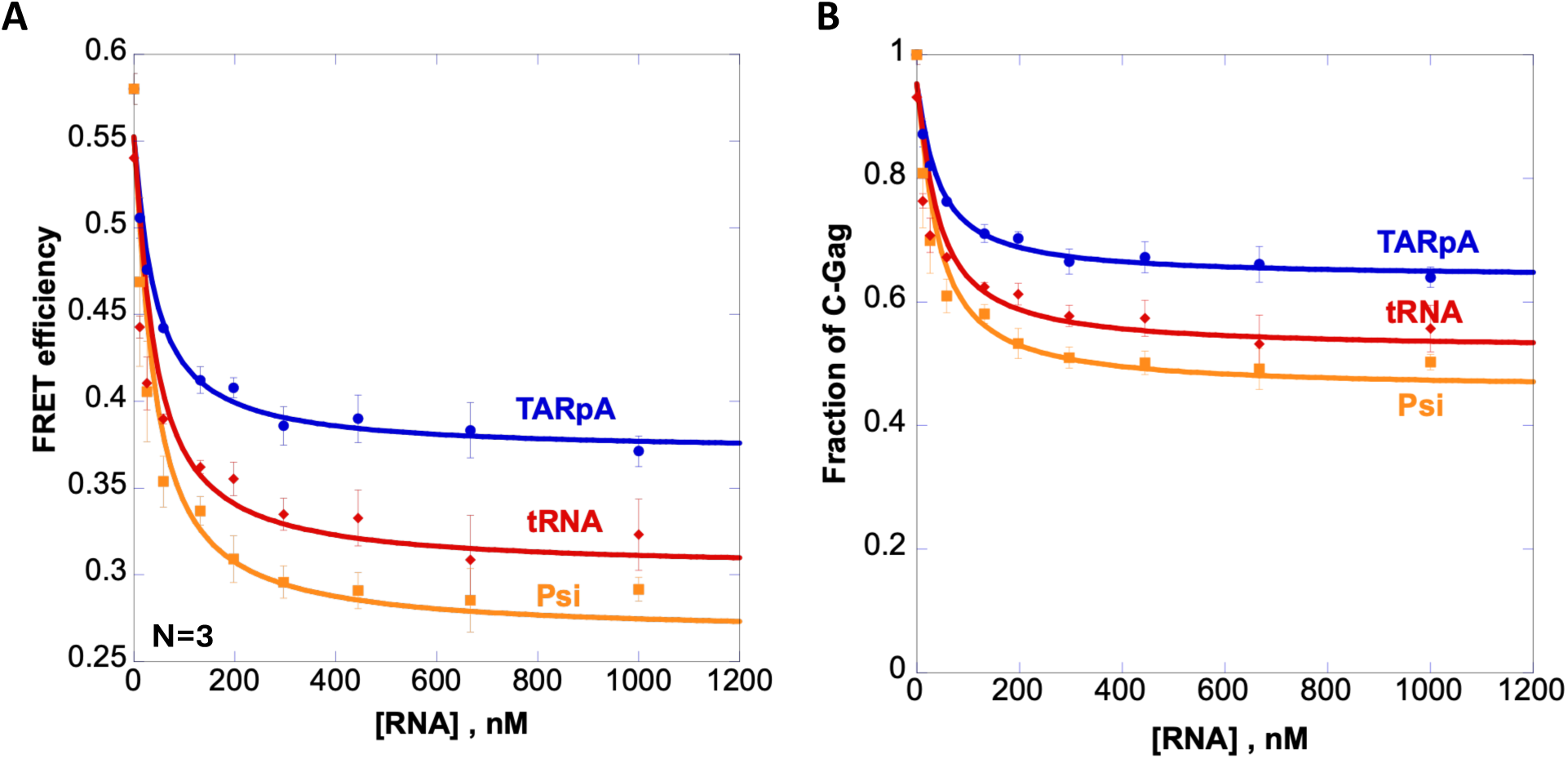
FRET efficiency change of dual-labeled WT Gag induced by RNA binding at 100 mM NaCl. A) FRET value of 20 nM Gag vs [RNA] for TARpA (blue), tRNA (red), and Psi (orange) RNA. Curves are the fits of measured data points using Eqs. 5, 7, and 8. B) Fraction of C-Gag as a function of [RNA] calculated from data in panel A according to Eq. 2.

The concentration at which 50% change in FE occurs was similar for all RNAs (∼20 nM). This value is similar to the Gag concentration used in these experiments, suggesting that the Gag-RNA dissociation constant is too small to be measured under these conditions, i.e., *K_d_* < [*Gag*] = 20 nM. In this case, we fit the Gag-RNA titration data as described by eq. 8 with the fraction of RNA-bound Gag described by eq. 6. Based on fits of the data shown in Figure 3 (solid lines), we estimated the fraction of C-Gag upon binding to a particular RNA molecule, *θ_C_^R^*, and the corresponding C-to-E transition free energy, *θ_CE_^R^*, (calculated according to eq. 4). As summarized in Table 1, the fraction of C-Gag decreases from 0.95 in the absence of RNA (at 100 mM NaCl) to *θ_C_^R^* ∼ 0.65, ∼0.54, and ∼0.44 upon saturated binding of TARpA, tRNA, and Psi, respectively. The C-to-E transition free energy decreases from *g_CE_^0^* ≈ 2.95 in the absence of RNA to *g_CE_^R^* ∼ 0.62, 0.16, and − 0.08 for TARpA, tRNA, and Psi, respectively. Thus, Gag binding to all RNAs tested resulted in a decrease in the C-to-E transition free energy by ∼2 − 3 *k_B_T*. The difference in the transition free energies determined for Gag bound Psi versus TARpA (δg_CE_^R^ ∼0.7) suggests that Psi-bound Gag is 1.4-fold more likely to be extended than TARpA-bound Gag.

**Table 1.** Fraction of compact Gag, free energy of the C-to-E transition of monomeric Gag, and Gag-RNA dissociation constant under different solution conditions.

| Protein and RNA | $\theta_C^a$ | $g_{CE}, k_B T^b$ | $K_d^c, \text{nM}$ | $K_d^c, \text{nM}$<br>+ 0.3 mM IP6 | $K_d^c, \text{nM}$<br>+ 20 nM tRNA<sup>Lys</sup> |
| --- | --- | --- | --- | --- | --- |
| Gag alone | 0.95 | 2.95 | — | — | — |
| Gag + TARpA | 0.65 | 0.62 | <20 | 162 | 500 |
| Gag + tRNA<sup>Lys</sup> | 0.54 | 0.16 | <20 | 120 | — |
| Gag + Psi | 0.44 | -0.08 | <20 | 36 | 76 |

<sup>a</sup> Fraction of C-Gag, $\theta_C$ ; <sup>b</sup> Free energy of C-to-E transition, $g_{CE}$ , in units of $k_B T$ ( $1 k_B T = 0.59 \text{ Kcal/mol}$ ), where $k_B$ is Boltzmann's constant in J/K and T is the absolute temperature in kelvin (K); <sup>c</sup> Gag-RNA dissociation constant in the absence or presence of inositol hexakisphosphate (IP6) or tRNA<sup>Lys</sup>. All measurements were performed with 20 nM Gag in 100 mM NaCl.

### IP6 alone partially destabilizes C-Gag

We next characterized the effect of the abundant cellular assembly co-factor, IP6, on Gag conformation. Titration of IP6 into a solution of 20 nM Gag led to an ∼0.13 reduction in FE that saturated at ∼20 μM IP6 (see Figure 4A). This corresponds to a decrease in the fraction of C-Gag from ∼0.95 to 0.74 and a corresponding C-to-E transition free energy decrease from ∼3 *k_B_T* for Gag alone to ∼0.8 *k_B_T* upon saturated IP6 binding (Table 1).

**Figure 4.**
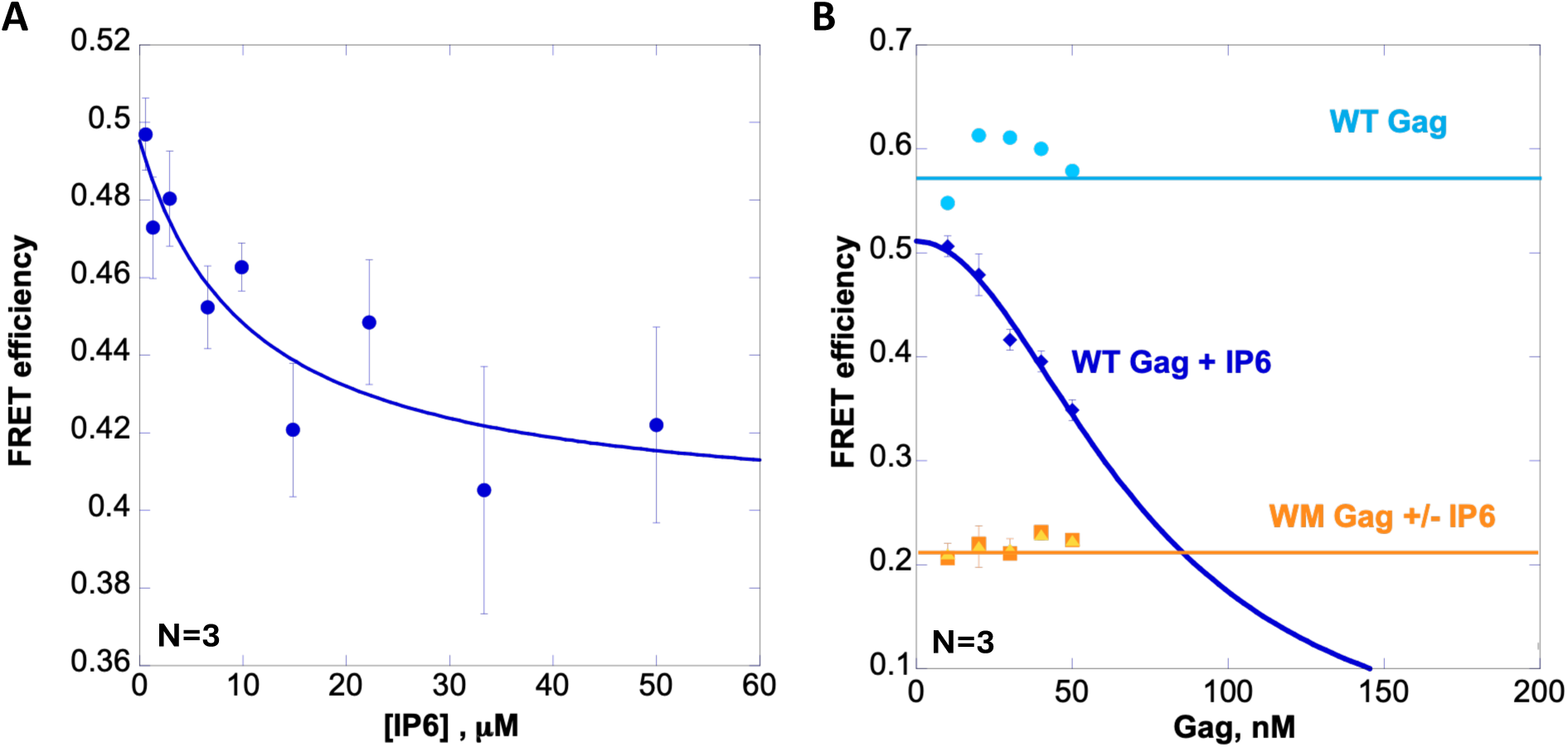
Effect of IP6 on Gag conformation. A) FRET efficiency as a function of IP6 for dual-labeled Gag (20 nM). The curve is drawn to guide the eye and is not fitted to the data. B) FRET efficiency as a function of dual-labeled WT Gag and WM-Gag concentration (10-50 nM) in the absence (lighter blue/orange) and presence (darker blue/orange) of IP6 (0.3 mM). The dark blue curve is a fit of the data to Eq. 9 with K_dim_ = 73±4 nM.

### Gag-RNA titration in the presence of IP6 reveals Gag binding differences between RNA molecules

We next performed Gag-RNA titration experiments in the presence of 0.3 mM IP6. Although the FE_max_ values were lower than in the absence of IP6 (as expected due to the IP6 effect on conformation) the observed decreases in FE as a function of RNA concentration plateaued at a value similar to that observed in the absence of IP6 (see Figure S3 for ±IP6 RNA titration comparison). However, in the presence of IP6, different concentrations of each of the RNAs tested were required to observe the maximum effect on FE. In this case, we fit the data in Figure 5 to the Gag/RNA binding isotherm given by eq. S7. Fitted K_d_ values for Gag-RNA binding in the presence of 0.3 mM IP6 were 162 nM, 120 nM, and 36 nM for TARpA, tRNA, and Psi, respectively (Table 1). Thus, competition with 0.3 mM IP6 weakens Gag-RNA binding, and under these conditions ∼4-fold stronger Gag binding is observed to Psi compared to TARpA.

**Figure 5.**
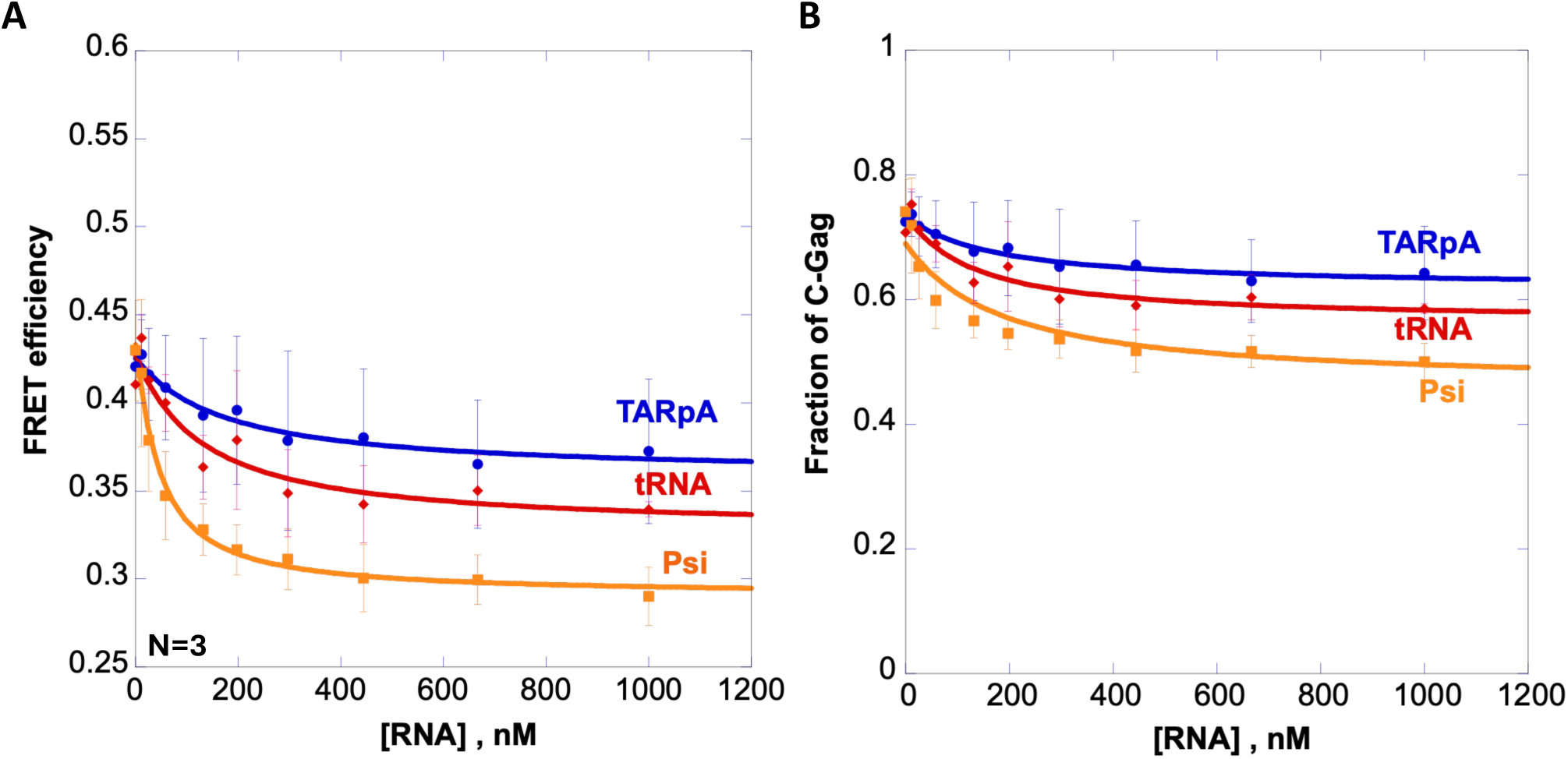
FRET efficiency change of dual-labeled WT Gag induced by 0.3 mM IP6 and RNA binding at 100 mM NaCl. A) FRET value of 20 nM Gag vs [RNA] for TARpA (blue), tRNA (red), and Psi (orange) RNA. Curves are the fits of measured data points using Eqs. 5, 7, and 8. B) Fraction of C-Gag as a function of [RNA] calculated from data in panel A according to Eq. 2.

### Psi competes with tRNA for Gag binding more effectively than TARpA

We next preincubated 20 nM Gag with tRNA^Lys3^ in a 1:1 ratio prior to RNA titrations with Psi and TARpA to mimic cellular conditions, in which the MA domain of Gag binds tRNA before membrane association (Kutluay et al. 2014; Thornhill et al. 2020). A decrease in FRET was observed after tRNA preincubation, consistent with the results of tRNA titrations (see Figure 3). Saturating concentrations of both Psi and TARpA RNAs induced FE decreases similar to those observed in the absence of tRNA (Figure 6). However, the effective K_d_s measured under these conditions are 500 nM and 76 nM for TARpA and Psi, respectively. Thus, tRNA is an even stronger competitor for Gag binding than IP6; under these competitive binding conditions the difference in the TARpA and Psi RNA binding affinities is ∼6-fold. (Table 1).

**Figure 6.**
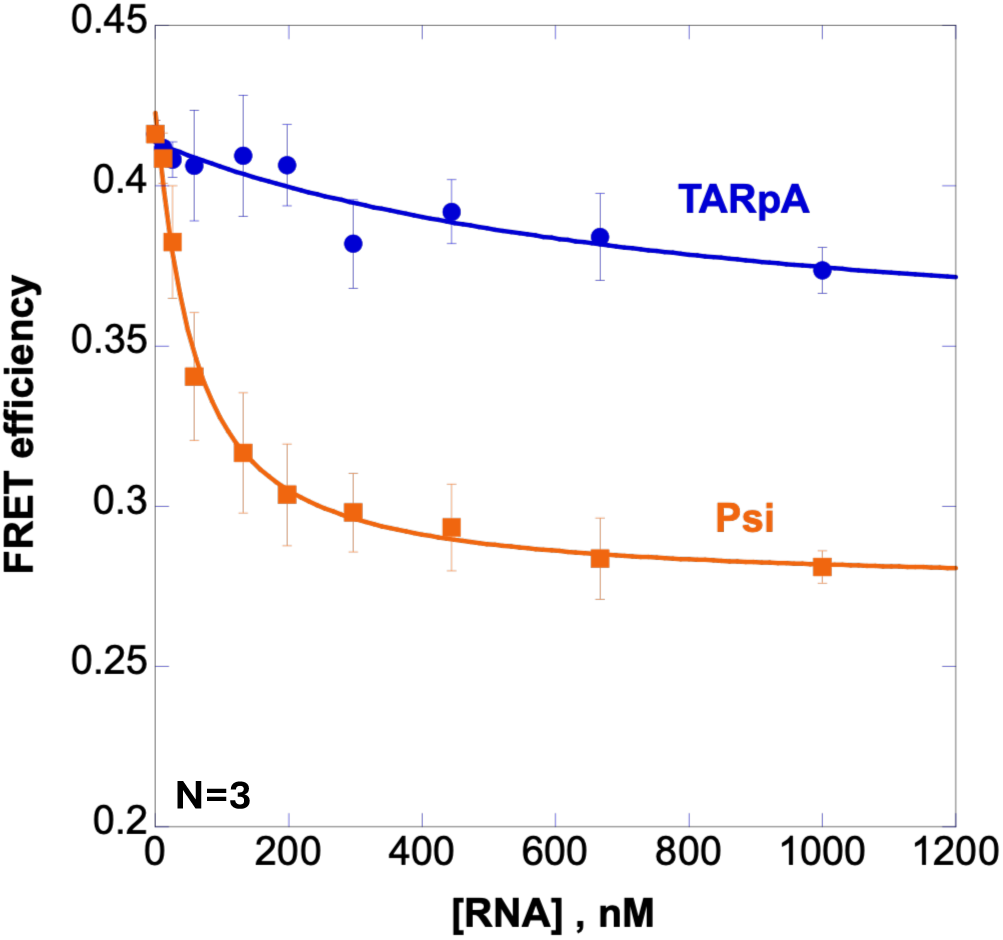
FRET efficiency change of dual-labeled Gag preincubated with tRNA^Lys3^ as a function of RNA concentration. WT Gag (20 nM) was preincubated with 20 nM tRNA^Lys^ and titrated with TARpA (blue) or Psi (orange). The curves are fits using Eqs. 5, 7, and 8.

### Higher Gag concentrations promote dimerization – induced Gag extension in the presence of IP6

The FE of Gag was next measured as a function of Gag concentration at constant IP6 (0.3 mM) in the absence of RNA. The FE value decreased with increasing Gag concentrations in the presence of IP6, but not in its absence (Figure 4B). For technical reasons, Gag concentrations were limited to 50 nM in these experiments. Assuming that each dimerized Gag is extended with FRET = 0, we can fit our FE data to Eq. 9, which yields an apparent Gag dimerization constant of K_dim_ = 73±4 nM in the presence of 0.3 mM IP6. This value is well below the reported dimerization K_d_ of 5-10 µM measured in the absence of the assembly co-factor (Datta et al. 2007b). To provide further evidence that this transition reflects Gag dimerization, we repeated the titration using dual-labeled WM-Gag, which was shown to dimerize ∼100-fold weaker than WT Gag (Datta et al. 2007b). No change in conformation was observed for WM-Gag either in the presence or absence of IP6 (Figure 4B, yellow and orange data points). Based on these results, as well as on the fact that WM-Gag is largely in the E-state in the absence of other factors (Figure 2A), we conclude that the WM site participates in intramolecular C-Gag stabilization as well as in Gag-Gag dimerization, and WM mutation leads to the loss of both interactions.

### Psi facilitates Gag oligomerization more effectively than TARpA

We next used MP to directly probe the effect of specific RNAs on Gag oligomerization. The molecular masses of Gag, TARpA, and Psi are 56.6 kDa, 35.7 kDa, and 36.4 kDa, respectively. In the presence of Gag and TARpA, two complexes were observed corresponding to 1:1 and 2:1 TARpA:Gag (Figure 7A). Psi binding to Gag resulted in five major complexes with up to four Gag molecules bound to a Psi dimer (Figure 7B). Although a small proportion (< 0.1) of TARpA had two Gag molecules bound, the majority interacted with only one Gag molecule (Figure 7C). Psi significantly promoted Gag oligomerization, with the RNA dimer recruiting up to five Gags and complexes containing more than one Gag molecule occupying over 50% of the total counts (Figure 7D). This difference between TARpA and Psi was not eliminated by the WM mutation; Psi still facilitated higher-order WM-Gag complex formation, although the overall fraction of Gag oligomers was reduced by a few-fold (Figure 7B,D). As for WT Gag, TARpA complexes primarily involved binding of a single WM-Gag.

**Figure 7.**
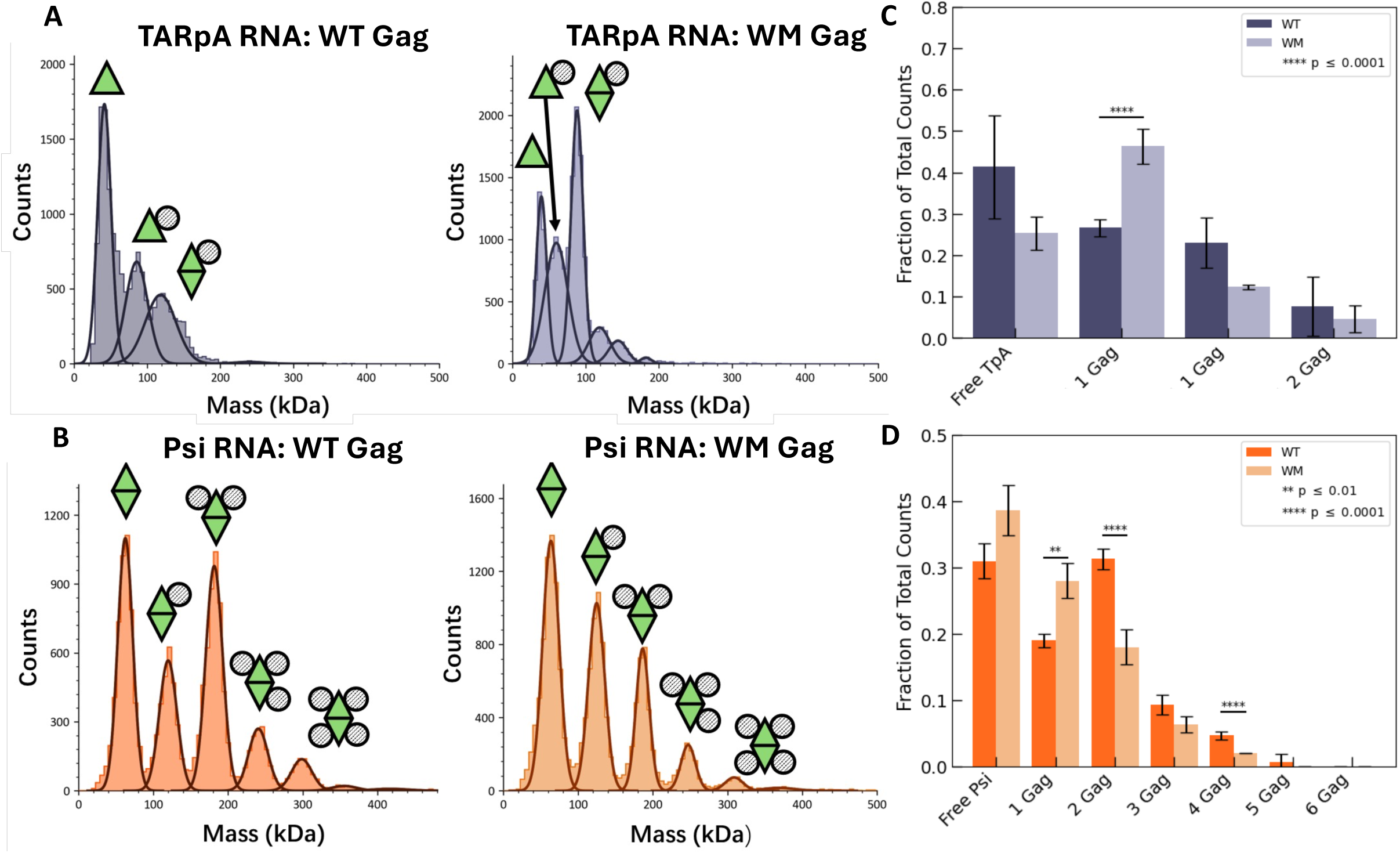
Mass photometry histograms. (average of three replicates) comparing A) TARpA RNA and B) Psi RNA binding with WT Gag (left) or WM-Gag (right). The green triangle represents one RNA molecule and the grey circle represents Gag. Summary of the fraction of complexes with different Gag stoichiometry for TARpA RNA (C) and Psi RNA (D).

### Atomistic MD simulations capture the enhanced flexibility of Gag monomers in the presence of Psi RNA-NC specific binding

All-atom MD (AAMD) simulations were carried out to study the impact of Psi RNA binding on Gag conformation and dynamics. We first monitored the end-to-end distance (R_ee_) of Gag in two sets of simulations: (1) (-)RNA: myristoylated Gag monomer (Gag monomers) alone in solution, (2) (+)Psi RNA: Psi RNA bound to Gag monomers in solution. R_ee_ is the distance between the center-of-mass (COM) of the N-terminal 10 residues of the MA domain and the COM of the C-terminal 10 residues of the p6 domain. All simulations started with the same initial extended structure of Gag monomers with R_ee_ ∼12 nm (Figure 8A,C). In the (-)RNA simulations, all time traces converged to a compact Gag monomers structure with R_ee_ values of ∼6 nm at the end of the trajectories (Figure 8A,B). This result is consistent with our experimental observation that the majority of Gag molecules are in the C-state in the absence of high salt, RNA or IP6 (see Figure 2, Table 1). In contrast, the (+)Psi RNA simulations revealed enhanced conformational flexibility of Gag monomers molecules (Figure 8C). Abrupt R_ee_ changes are observed, consistent with transitions between C-Gag (R_ee_ values as low as ∼4 nm) and E-Gag (R_ee_ values near 10 nm).

**Figure 8:**
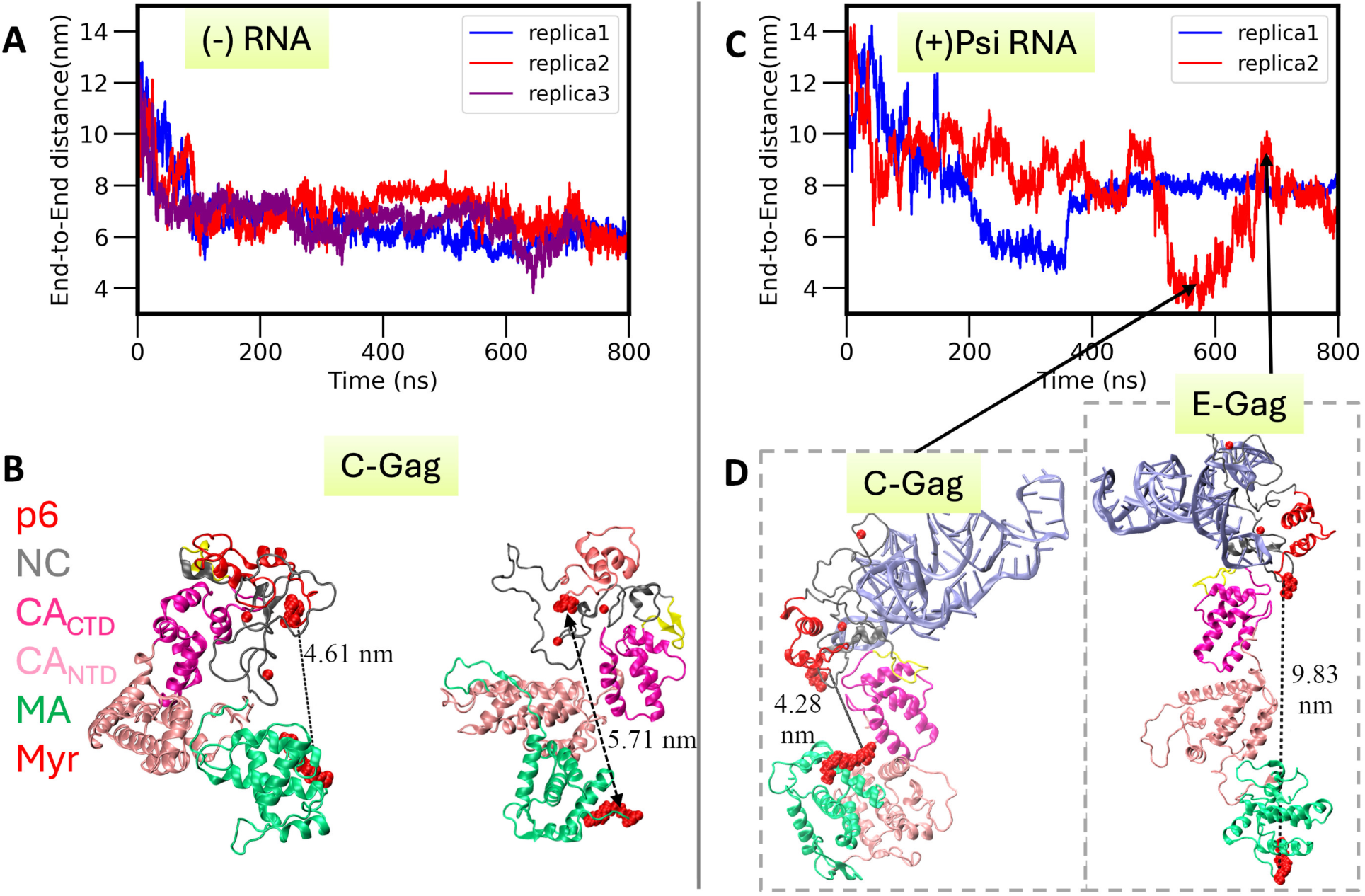
Conformational change of Gag polyprotein measured by monitoring end-to-end distance (R_ee_) using all-atom MD simulations. (A,C) Time-series of R_ee_ for MD trajectories of Gag monomers in the absence ((-)RNA) and presence of Psi RNA-NC specific binding ((+)PsiRNA). (B,D) Snapshots of Gag monomers in compact (C-Gag) conformations extracted from (-)RNA trajectories (B) and both C-Gag and extended (E-Gag) conformations in the (+)Psi RNA system (D). Black arrows point to the timeframes of the trajectory representative C-Gag and E-Gag structures are taken. Domains of Gag polyprotein are colored as defined in panel B. RNA is shown in ice blue in panel D. In panels B and D, N-terminal and C-terminal residues are shown in red and R_ee_ values are indicated.

### NC interdomain interactions stabilize C-Gag states in the absence of Psi RNA-NC binding

As observed from the AAMD simulations while monitoring end-to-end distance, Gag molecules can attain C-Gag conformations in both (-)RNA and (+)Psi RNA systems. However, conformational transitions to E-Gag occur only in the presence of Psi RNA-NC binding. The Gag MA domain is connected to the CA_NTD_ through a flexible linker. All of the AAMD trajectories in the absence of RNA sampled the C-Gag state with stable MA:CA_NTD_ binding (**Figure S4**A,C). Two out of three trajectories also captured the MA domain in close proximity to the CA_CTD_ (**Figure S4**B,D). However, no direct interactions or stable binding was observed between these two domains in a full-length Gag molecule. Stable MA:CA_CTD_ binding was predicted based on previous simulations of separate MA and CA_CTD_ domains (Lin et al. 2019).

To investigate inter-domain interactions mediated by the NC domain in Gag, we measured the minimum distance between MA:NC and CA_NTD_:NC domains in the (-)RNA and (+)Psi RNA AAMD systems. In the absence of RNA, we observed stable NC-CA_CTD_ and NC-CA_NTD_ binding for all replicas of the AAMD simulations, driven mainly by electrostatic interactions (Figure 9A, Figure S5A, B). One out of three trajectories (Figure 9A, trajectory 2 shown in Figure S5C) also sampled direct binding of the NC domain acidic residues with the lysine-rich domain of MA. In the (+)Psi RNA condition, no stable interactions of the NC domain with either CA_NTD_ or MA were captured in the simulations (Figure 9B, D), and the basic residues of NC were observed to primarily bind RNA.

**Figure 9:**
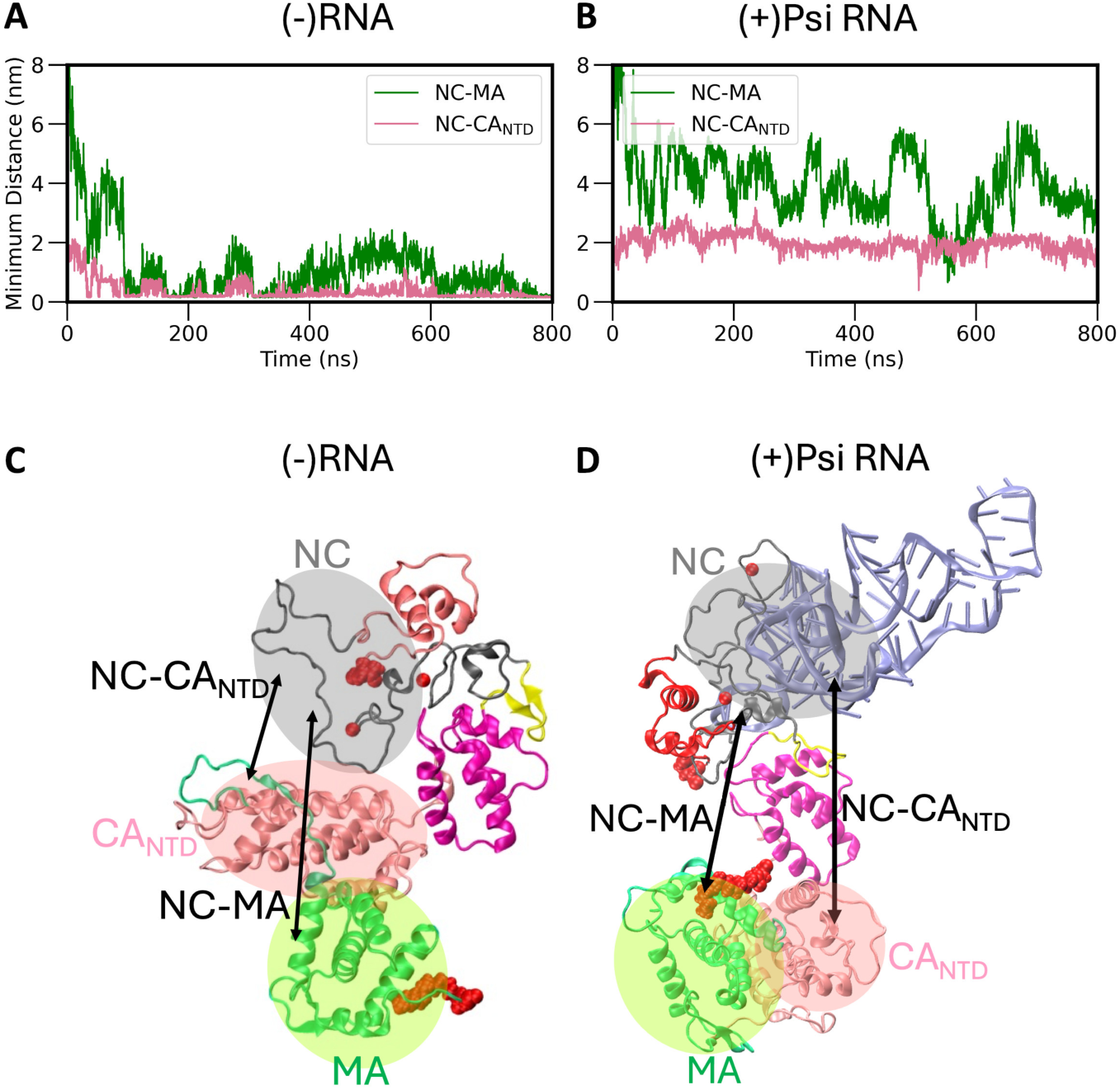
Inter-domain Gag interactions in the presence and absence of Psi RNA-NC specific binding. (A-B) Minimum distance from the MA and CA_NTD_ domains to the NC domain in the absence (-) and presence (+) of Psi RNA. (C-D) Representative conformations of C-Gag. Arrows in panels C, D indicate the distances measured.

### Molecular mechanism of the C-Gag to E-Gag transition

In long-timescale (+)Psi RNA unbiased AAMD simulations, Gag monomers polyprotein was found to exhibit spontaneous transitions from compact to extended states (Figure 10A). We analyzed several structural parameters to better understand the dynamics of this conformational change. As mentioned earlier, the structural domains of the Gag molecule are connected by disordered linker regions. To monitor CA flexibility, which is regulated by the linker between the CA_NTD_ and CA_CTD_, we measured the angle between Glu244 (CA_NTD_), Gln307 and Thr347 (both in the CA_CTD_). This angle showed a sharp rise from 80^0^ to 150^0^ during the simulation time window plotted to the left of the dotted line in Figure 10B. This conformational change is correlated with MA:CA_NTD_ unbinding dynamics (Figure 10C). To monitor the latter, we measured the number of contacts between MA domain residues 2-110 and CA_NTD_ residues 147-262, with a cut-off distance of 4.5 nm. The ∼ 500 contacts, observed in the compact form of Gag at the start of the AAMD simulation, decrease to ∼ 300 at 650 ns and fall sharply to 0 at ∼ 660 ns. During this last window (660-685 ns), the end-to-end distance exhibits a transition from ∼6 nm to ∼10 nm (Figure 10A).

**Figure 10:**
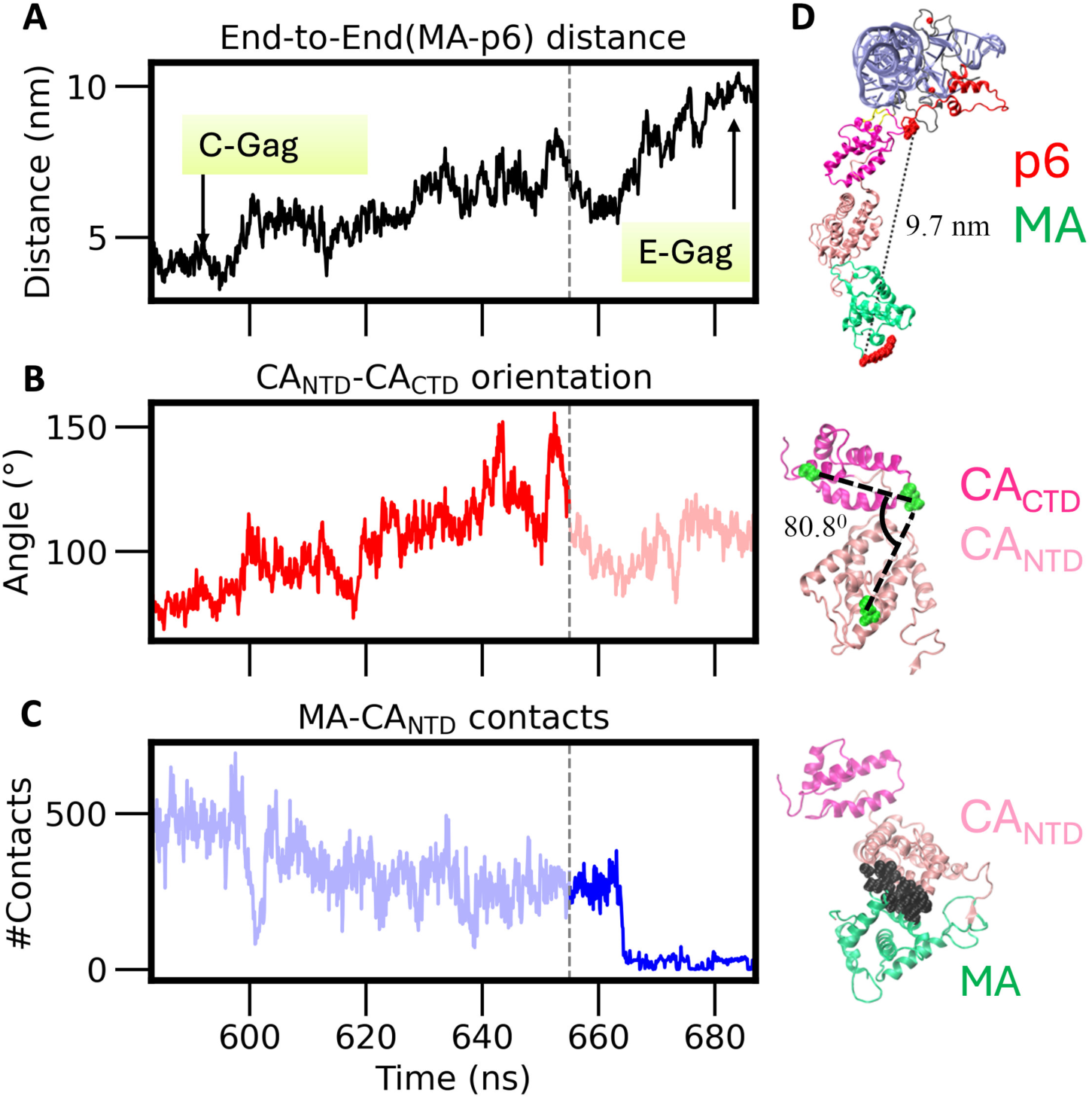
Mechanism of C-Gag to E-Gag transition of Gag monomer in solution in the presence of Psi RNA-NC specific binding. Time-series of MA-p6 end-to-end distance (A), CA_NTD_:CA_CTD_ domain orientations (B), and MA domain unbinding from the CA_NTD_ (C). Measured structural parameters in the left panels, with their magnitudes in the representative structures are shown in (D). The vertical dotted line separates the conformational transition into two time windows, where the structural changes shown in (B) and (C) facilitate progression from the compact C-Gag to the extended E-Gag conformation.

## Discussion

Using dual-labeled Gag and an ensemble FRET approach, we demonstrated that the C state of Gag is stabilized in part by electrostatic interactions, which aligns with previous MD simulation results showing that MA interacts with CA_CTD_ through multiple charged as well as polar residues (Lin et al. 2019). Interestingly, the FRET efficiency of free WM Gag (0.22) was much lower than that of WT Gag (0.55), suggesting that the WM site in the CA_CTD_ is not only involved in intermolecular Gag dimerization, but contributes to stabilizing C-Gag through intramolecular contacts. These results imply that the intra-Gag contacts stabilizing C-Gag have a hydrophobic component involving the WM site, as well as electrostatic contributions predicted previously by MD simulations (Lin et al. 2019) and in the present work. The observed contribution of the WM site to C-Gag stabilization is in agreement with recent studies in cells, which reported a significant reduction in the cytoplasmic C-Gag fraction of the WM mutant compared to WT Gag (Zeiger et al. 2024).

RNA binding induces a conformational shift of WT Gag from C-Gag to E-Gag both in the absence and presence of IP6. In contrast, previous single molecule studies that tested the effect of polyA_25_ DNA binding on Gag conformation reported that no change was observed in the absence of IP6 (Munro et al. 2014). All RNAs tested here induced a conformational shift but the effect of Psi RNA on the average FRET efficiency change was greater than that of non-Psi RNAs (Figure 11). We hypothesize that the specific interactions between Psi RNA and the NC domain are optimized by formation of the E state. IP6 is a critical host cell assembly factor known to bind to CA in both the immature and mature particles (Dick et al. 2018; Gupta et al. 2023). Based on its high negative charge, IP6 is also capable of binding to the basic domains of MA and NC (Datta et al. 2007b). Even in the absence of RNAs, IP6 binding induced a conformational change toward E-Gag in our experimental studies. Upon RNA titration, the minimum FRET of WT Gag reached a similar plateau as for experiments performed in the absence of IP6. Without this co-factor, Gag binding to all three RNA molecules tested in 100 mM NaCl was too strong to be measured (K_d_ <20 nM). However, the K_d_ values measured for the three RNAs in the presence of IP6 were significantly different (36 nM for Psi, 120 nM for tRNA, and 162 nM for TARpA) suggesting that IP6 and RNAs bind to Gag at the same sites and Psi is the most effective competitor. Thus, in the presence of IP6, Gag can more effectively discriminate between Psi and non-Psi RNAs.

**Figure 11.**
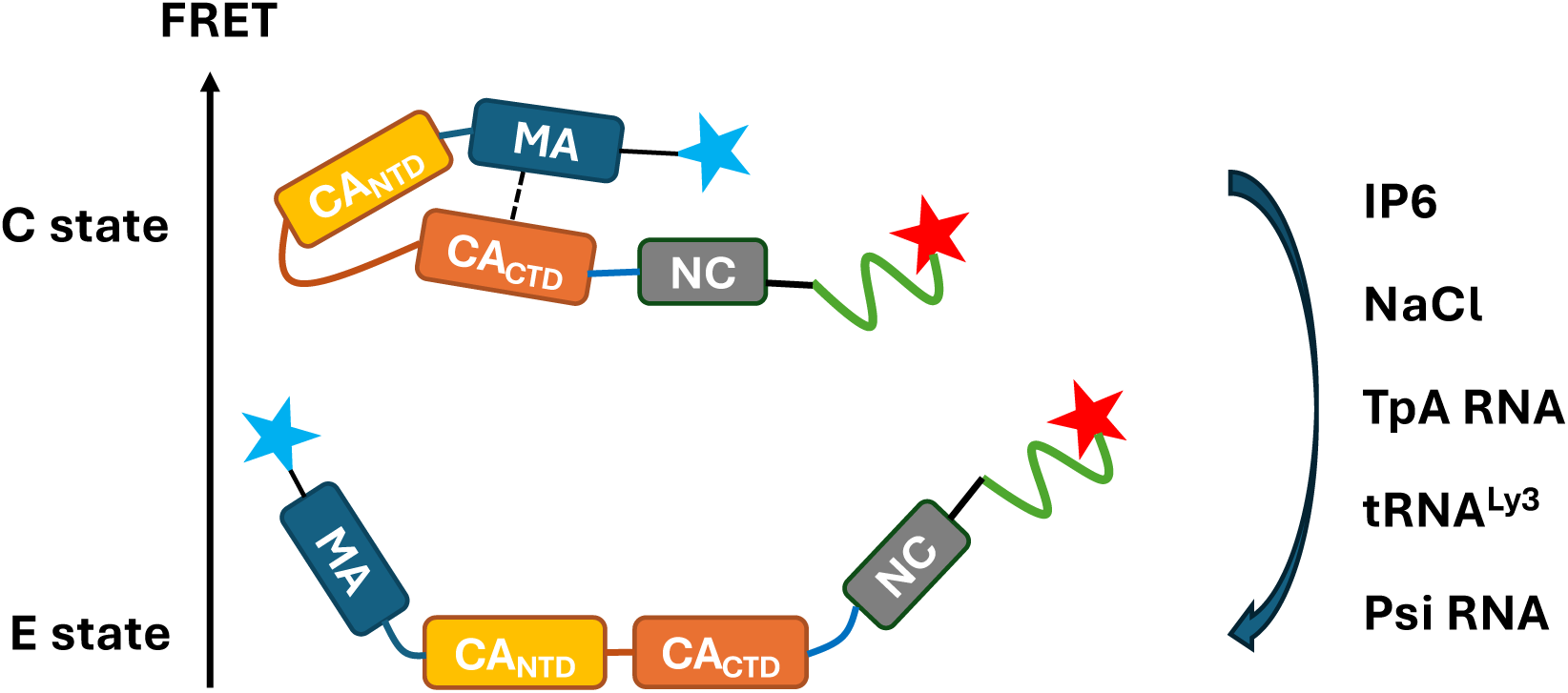
Scheme showing factors that impact Gag conformational change. Two major states with different FRET efficiencies are shown: C state for compact Gag and E state for extended Gag. While Gag is mostly in the C state at low salt and in the absence of other factors, the curved arrow indicates the conformational equilibrium shift induced by different factors in the order of the strength of their effect from weak to strong (top to bottom).

The K_dim_ value measured for WT Gag dimerization in the absence of RNA in high salt (0.5 M NaCl) solution conditions was ∼5-10 µM (Datta et al. 2007b). This value increased to ∼500 µM for WM-Gag. In our studies, the FRET efficiency at various concentrations of Gag revealed a sharp decrease with increasing WT Gag (+ IP6). Since Gag dimerization via CA-CA interactions is expected to promote Gag extension, the FRET decrease observed in the presence of IP6 likely reflects dimerization with an estimated K_dim_ ∼70 nM. Previous biophysical studies showed that IP6 promoted a Gag monomer-trimer equilibrium rather than the monomer-dimer equilibrium observed in the absence of the assembly co-factor (Datta, 2007b). Thus, our data may reflect higher order multimerization in the presence of IP6. The high negative charge of IP6 likely neutralizes the basic residues in the MA and NC domains, which cause repulsion when Gag proteins oligomerize. Gag was observed to dimerize at concentrations of ∼500 nM in cells (Fogarty et al. 2014; Hendrix et al. 2015) and *in vitro* studies performed in the presence of a short (GT)_3_ DNA oligonucleotide in the absence of IP6, observed dimerization through an NC interface at concentrations as low as ∼100 nM Gag (Zhao et al. 2019), a value similar to our K_dim_. Our results suggest that Gag-Gag interactions are very sensitive to the presence of RNA, IP6 and salt, all of which induce conformational changes. Dimerization is strongest in the absence of RNA when charge neutralization and IP6-induced extension of Gag promote intermolecular interactions.

MP binding data also support the conclusion that the conformational change of Gag induced by specific RNA binding contributes to Gag oligomerization. Psi RNA promoted the higher-order oligomerization of Gag, whereas TARpA RNA failed to do so. WM-Gag similarly formed complexes with two to four Gag molecules in the presence of Psi but not TARpA, which indicates that Psi facilitates complex formation by affecting Gag conformation and by promoting a higher local concentration of Gag with multiple proximal NC binding sites.

The present study is the first to measure the influence of specific viral and non-viral RNAs on Gag conformation, with important implications for specific HIV-1 gRNA packaging and virion assembly. The results support a two-state model for major conformational states, with the C state stabilized by a combination of electrostatic and hydrophobic interactions (Munro et al. 2014; Su et al. 2018; Lin et al. 2019). The factors driving selective packaging of viral gRNA into virions have been actively investigated for decades. *In vitro* studies consistently showed that most RNAs bind to Gag with similar affinity under physiological conditions. In this work, the combination of IP6 and specific Psi RNA binding showed the most significant impact on Gag conformational change and dimerization propensity. The MA domain of Gag shows a tRNA-binding preference in cells (Kutluay et al. 2014). Mimicking these conditions by pre-incubating Gag with tRNA prior to Psi and non-Psi RNA titration, provided additional support for the specific effect of Psi RNA on Gag conformational change.

To complement the experimental results, AAMD simulations investigated the impact of NC-Psi RNA specific binding on the conformational dynamics of monomeric Gag in the solution. The conformational transitions of multidomain proteins are multistep processes that include inter-domain open-to-close hinge motions as well as folding/unfolding events of structural domains. Because large-scale conformational transitions of HIV-1 Gag are rare on accessible simulation timescales, characterizing them at the molecular level is difficult. Our long-timescale AAMD simulations captured the transition of compact to extended state of monomeric Gag (C-Gag to E-Gag) in the presence of NC-Psi RNA specific binding, unlike the simulations of free Gag monomer. Together, the simulated data explain the experimental observations that Psi RNA promotes the equilibrium shift towards E-Gag from C-Gag more efficiently than non-Psi RNAs due to NC:Psi RNA specific binding. Our previous MD studies identified the role of RNA in modulating the dynamics of the MA-CA_NTD_ linker in the context of membrane-bound Gag multimer conformational transition (Banerjee and Voth 2024). In the current study, our simulations elucidate the mechanism of the Psi RNA-induced conformational transition of monomeric solution-phase Gag, which is driven by CA_NTD-CTD_ orientational flexibility and MA-CA_NTD_ unbinding dynamics.

Recent experiments in HIV-1-transfected cells showed that expressing Gag mutants with reduced amounts of cytoplasmic C-Gag correlated with infectivity defects (Zeiger et al. 2024). This result suggests that the stability of C-Gag may play a physiological role in the HIV-1 lifecycle. The present work allows us to estimate the preference of Gag dimer formation on Psi. If E-state occupancy independently contributes to dimer formation by each Gag monomer, the ∼1.4-fold higher E-state probability of Psi-bound Gag versus TARpA-bound Gag would be expected to produce an approximately two-fold (1.4^2^) increase in E-Gag dimer formation. In addition, the ∼6-fold stronger E-Gag binding to specific sites on Psi vs TARpA (following pre-incubation with tRNA), leads to an estimated 36-fold (6^2^) higher probability of dimer formation on Psi. As TARpA is known to be one of the least specific Gag binders (Webb et al. 2013), together these contributions suggest an estimated upper-limit enhancement of Gag dimer formation on Psi relative to TARpA/non-Psi RNA of ∼ 72-fold. Recent RNA-seq measurements suggest that total HIV-1 RNA represents ∼1% of the mapped mRNA reads in the cell at 16-24 h post infection (Kibe et al. 2025), suggesting that factors other than specific Gag oligomerization on Psi likely play a role in selective unspliced gRNA packaging. The existence and stability of the C-Gag conformation may have other functions in the HIV-1 lifecycle. For example, C-Gag stability may moderate the strength of Gag-Gag interactions within the immature lattice or affect the rate of virion assembly. Future studies may uncover yet unappreciated roles of intra-Gag interactions.

In summary, the integrated experimental and computational approaches described in this work establish the key role of NC-Psi RNA binding in the conformational switch of Gag from a compact molecule to the assembly-competent extended state. The abundant cellular factors, IP6 and tRNA, are known to bind to cytoplasmic Gag in the initial phases of the assembly process prior to plasma membrane binding. We show here that these factors also promote Gag extension as well as oligomerization and specific Psi RNA binding. Overall, this study reveals how the highly flexible multi-domain Gag polyprotein leverages both viral and host cell factors to sample and stabilize distinct conformations, thereby orchestrating the tightly regulated progression of viral assembly.

## Materials and Methods

### Protein induction and purification

A full-length Gag construct for dual labeling was designed based on the plasmid of Gag-MBP-His received as a gift from David Millar (Scripps Research). Two additional glycine residues were inserted after the start codon of the Gag sequence to form an N-terminal triglycine motif recognized by Sortase (Mao et al. 2004). Since multiple cysteine residues naturally exist in Gag, a KCK motif was inserted at the 494-496 position to enhance the reactivity of this specific cysteine at the C-terminal (Larson et al. 2017). The monomeric variant was generated from the WT Gag construct by mutating W316 and M317 to alanine. The primer sets used for Site-directed, Ligase-Independent Mutagenesis (Chiu et al. 2004) are listed in Table S1. After confirming the sequence, the plasmids were transformed into Rossetta cells for auto-induction. The cells were induced in ZY-5025M as described previously (Studier 2005). To inoculate 1 L of ZY-5025M, 100 ml of overnight culture was prepared in LB media by shaking at 37 °C. The incubation temperature was adjusted to 18 °C after the OD_600_ of the 1 L culture reached 2.0. The induction lasted at least 24 h and ensured a typical yield of 10 mg of Gag was obtained following the Gag preparation procedure described previously (Sarni et al. 2020). Briefly, the cell pellet was lysed by sonication and removed nucleic acid by polyethylenmine precipitation. The supernatant was recovered after centrifugation. Gag was precipitated by saturate ammonium sulfate and resuspended after collecting the pellet by centrifugation. The resuspended Gag solution was load to Ni^2+^ resin and the purified Gag was eluted from 50-300 mM imidazole buffer. The fractions of Gag were collected after gel checking and added 3 mg TEV protease per 20 mg of Gag to remove MBP-His tag. Uncleaved Gag, MBP-His tag, and TEV protease were removed by Ni^2+^ resin. The fractions of cleaved Gag were buffer exchanged to salt buffer for Heparin Column (5 mL, Cytiva). The purified Gag was eluted from 600-900 mM NaCl buffer. The fractions of Gag were collected after gel checking and concentrated to ∼10 ml by Vivaspin 10 kDa (Cytiva). The concentrated Gag was dialyzed to Gag storage buffer (20 mM Tris-HCl pH 7.5, 5 mM βME, 500 mM NaCl, 1 µM ZnCl_2_, 10% glycerol). The Gag concentration was determined by measuring the A_280_ and using an extinction coefficient of 64400 M^-1^cm^-1^.

A His-tagged Sortase used for N-terminal labeling was produced in our lab using a published protocol (Dillard et al. 2019). The plasmid encoding 5M SrtA was purchased from Addgene (#51140) and transformed into Rossetta cells. To inoculate 1 L of LB, 100 ml of overnight culture was used, and 0.5 mM IPTG was added after the OD_600_ reached 0.6. The induction was performed overnight at 18 °C shaking. Briefly, the cell pellet was lysed by sonication and the supernatant was recovered by centrifugation. The supernatant was loaded onto Ni^2+^ resin and Sortase was eluted from 500 mM imidazole buffer. The fractions of Sortase were collected after gel checking and concentrated to ∼500 µM by Amicon 3 kDa (Sigma). The final product was dialyzed into 50 mM Tris-HCl pH 8 containing 150 mM NaCl and 10% glycerol. The Sortase concentration was determined by measuring the A_280_ and using an extinction coefficient 14440 M^-1^cm^-1^. All proteins were aliquoted and stored at-80 °C.

### RNA preparation

TARpA, Psi and tRNA^Lys3^ (sequences listed in Table S2) were generated by *in vitro* transcription with T7 RNA polymerase using Fok1-digested plasmids as the template. Transcription was carried out as described (Milligan et al. 1987). TARpA and Psi used the plasmid pMSMΔEnv containing HIV-1 NL4-3 cDNA as previously described (Webb et al. 2013). tRNA^Lys3^ used the plasmid pLysf119 (Shiba et al. 1997). Plasmid were purified from 0.5 L of overnight culture of DH5α cells with a Maxiprep kit (QIAGEN). The *in vitro* transcription reaction conditions were 1x TB buffer (80 mM HEPES-Na^+^, pH 8.0, 30 mM Mg(OAc)_2_, 10 mM DTT, 5 mM spermidine, 0.01% Triton-X-100), 4 mM NTPs, 0.1 µg/µl template, 1 U pyrophosphatase, and 25 µl/ml homemade T7 RNA polymerase. After a 4-hour incubation at 37 °C, the reaction was quenched by 50 mM EDTA and mixed with RNA denaturing dye (47.5% formamide, 0.01% SDS, 0.01% bromophenol blue, 0.005% xylene cyanol, 0.5 mM EDTA). The mixture was loaded onto an 8% 20×20 cm urea polyacrylamide gel and the desired RNA band was cut from the gel after running for 2 h at 400 volts. The RNA was extracted from the gel by shaking in elution buffer (0.5 M NH_4_OAc, 1 mM EDTA, pH 8.0) overnight at 37 °C. The solution was concentrated to about 400 µl by butanol extraction and then subjected to ethanol precipitation to obtain the RNA pellet. After resuspending the pellet in an appropriate amount of Milli-Q water, the concentration was determined by A_260_ measurement using a Nanodrop (Thermo Fisher) and the following extinction coefficients: 935693 M^-1^cm^-1^ (TARpA), 954383 M^-1^cm^-1^ (Psi), 664688 M^-1^cm^-1^ (tRNA^Lys3^).

### Dual-labeled Gag preparation

The purified Gag proteins (both WT and WM containing an N-terminal GlyGlyGly motif and a ^494^KCK^496^ motif) underwent the double-dye labeling protocol shown in Figure 1. Cy3 dye was appended to the N-terminal end through Sortase-mediated peptide labeling (Dillard et al. 2019). This enabled ligation between the triglycine motif at the N-terminus of Gag and the Cy3-LEPTGG peptide dissolved in 20 mM Tris-HCl pH 7.5 (purchased from Sigma-Aldrich). A typical large-scale reaction was performed in 100 µL by adding 10 mM CaCl_2_ and 5x Cy3-LPETGG peptide to 20 µM Gag (1x) in the Gag storage buffer (20 mM Tris-HCl pH 7.5, 500 mM NaCl, 5 mM βME, 1 µM ZnCl_2_). After a 1-hour incubation at 37 °C, the Cy3-Gag was buffer exchanged to a Cys labeling buffer (25 mM Tris-HCl pH 7.0, 500 mM NaCl, 100 µM TCEP, 25 µM ZnCl_2_) using a Zeba desalting column (Thermo Fisher) followed by Cy5-maleimide conjugation at the C-terminus. A dilution step was required for WT Gag concentrated to more than 10 µM to avoid protein aggregation. A typical reaction volume for this step was 200 µL with the Gag concentration at 10 µM. The six Cys residues in the zinc fingers of NC were protected by an additional ZnCl_2_ (50x the concentration of Gag) before the addition of 3x Cy5-maleimide (ThermoFisher). The dye stock was prepared by dissolving in DMSO and the concentration was determined using the extinction coefficient 250000 M^-1^cm^-1^. The reaction was quenched by 10x βME after a 2-min incubation at room temperature (RT). The dual-labeled Gag was then buffer exchanged into Gag storage buffer and the His-tagged Sortase was removed by Ni^2+^ resin before concentrating the final product by Vivaspin 10 kDa (Cytiva). The labeling efficiency of each dye was evaluated by SDS-PAGE and calculated by measuring the absorbance at A_280_ (Gag), A_555_ (Cy3), and A_648_ (Cy5) and the following extinction coefficients: 64400 M^-1^cm^-1^ (Gag), 15000 M^-1^cm^-1^ (Cy3), 25000 M^-1^cm^-1^ (Cy5). If the Gag concentration from A_280_ does not agree with the band intensity on gel, run 10 µl of final Gag product with a serial BSA standard (1-5 µg) on SDS-PAGE and calculate the Gag concentration by the BSA standard curve generated from the band intensity. A typical labeling stoichiometry was ∼55% for each dye. The final product was aliquoted and stored at-80 °C.

### Binding reaction setup and FRET measurement

All titration experiments were performed with 20 nM Gag in 20 mM HEPES pH 7.5, 50 mM NaCl, 1 mM βME, 2 mM Tris-HCl pH 7.5, 100 nM ZnCl_2_ (components from 10x diluted Gag storage buffer). For the NaCl titration, 37–500 mM NaCl to reach final NaCl concentration 87-550 mM. For the IP6 titration, 50 mM NaCl was added to reach final NaCl concentration 100 mM. A serial IP6 (pH 7.5) dilutions (5.8-500 µM) were prepared from 10.77 mM IP6 (pH 7.5) solution (10x dilution of 1.1 M stock by Milli-Q water + 4.2 µl of 4 M NaOH). 0.58–50 µM IP6 pH 7.5 was added to the final reaction. For the RNA titration, 50 mM NaCl (100 mM in final reaction), 1 mM MgCl_2_ and 11–1000 nM RNA was added. The measurements were taken after a 30-min RT incubation to reach equilibrium. For the tRNA preincubation test, a 10x concentrated Gag solution (200 nM) was incubated with 200 nM tRNA^Lys3^ at RT for 30 min before setting up the final reaction as the regular RNA titration. All RNAs were refolded at 10 µM in 50 mM HEPES, pH 7.5 before use. The steps included a 2-min incubation at 80 °C, a 2-min incubation at 60 °C, addition of 10 mM MgCl_2_, and an ice incubation for at least 30 min. A spectrofluorometer (FluoroMax-4, HORIBA) was used to measure the spectrums for FRET calculation. The direct Cy3 donor excitation spectrum (ex = 510 nm) was collected from 530 nm – 750 nm. The Cy5 acceptor excitation spectrum (ex = 610 nm) was collected from 630 nm – 750 nm.

### FRET calculation and model fitting

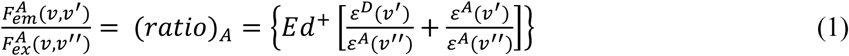

The FRET efficiency for each binding reaction was calculated using Eq. (1), the (*ratio*)*_A_* method (Clegg 1992). A MATLAB script was written for data analysis, and typical spectra are shown in Figure S1B. Briefly, to determine the quantity (*ratio*)*_A_*, we calculated the ratio of *F_em_^A^ (v,v’)* versus *F_ex_^A^ (v,v’)* from experimental measurements. *F_em_^A^ (v,v’)* is the acceptor emission at frequency *v* when the acceptor is excited through the FRET interaction (*v*′ excitation). First, a reference excitation spectrum of the donor only (D) was scaled to the donor’s maximum emission peak (570 nm for Cy3) in the experimental direct donor excitation spectrum (D+A). Once these spectra are aligned, the contributions to D+A that do not result from FRET can be obtained (Figure S1B, middle). Second, the quantity *F_em_^A^ (v,v’)* was obtained by subtracting the spectrum of D from the spectrum D+A resulting in the acceptor (Cy5) emission due to FRET (Figure S1B, bottom). *F_ex_^A^ (v,v’)* is the acceptor emission at frequency *v* when the acceptor is directly excited (*v*′′ excitation), obtained by Cy5 direct excitation. The FRET efficiency (E) was calculated from Eq. (1) using the experimentally determined quantity (*ratio*)*_A_* and the known values for the extinction coefficients *ε^A^(v’),ε^A^(v”),ε^D^(v’),ε^A^(v”)*, as well as d^+^, the labeling efficiency of the donor. For a fully labeled sample, d^+^ = 1 (100%). We used 0.55 due to the 55% Cy3 labeling efficiency of Gag. An advantage of this method is that the acceptor labeling efficiency (a^+^) does not need to be known as its contribution is cancelled in the ratio.

### Analysis of the C-Gag – to – E-Gag transition as a two-state system

Assuming that each Gag molecule is a two-state system(Munro et al. 2014) and normalizing to the maximum FRET value in low salt (see Figure 2), we can calculate the fraction of Gag molecules in C-state,

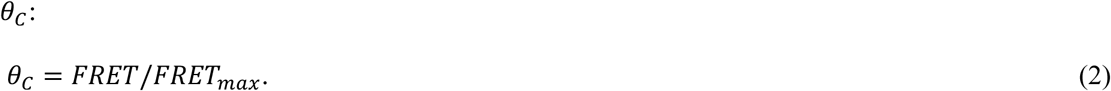

As for any two-state system, the fraction of C-Gag is related to the free energy of the C-to-E transition, *g_CE_*, as follows:

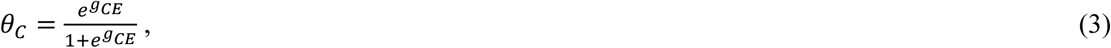

allowing for the estimate of *g_CE_*:

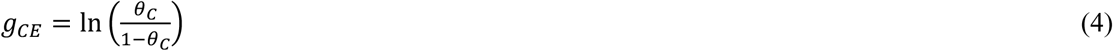

### Fitting of the Gag/RNA titration curves

We fit all of our titration curves using two assumptions: (i) the free energy of the C-to-E transition for the RNA-bound Gag molecule, *g_CE_^R^*, is lower than for Gag alone, *g_CE_^0^*; (ii) the fraction of RNA-bound Gag molecules, *f_R_*(*R*), can be described by the simple binding isotherm:

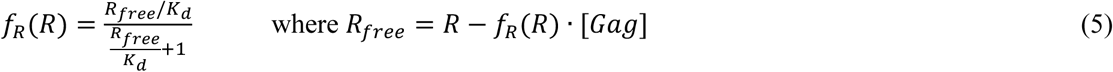

where *R* and *R_free_* are concentrations of the total (Gag bound and unbound RNA) and free RNA in solution, respectively. When binding is strong, such that *K_d_* < [*Gag*] = *G* all added RNA molecules bind Gag, and the fraction of RNA-bound Gag does not depend on the strength of binding (i.e., on *K_d_*):

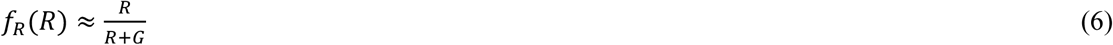

In contrast, when the concentration of interacting molecules, [*Gag*], *R* ≤ *K_d_*, eq. (5) can be used to find the explicit binding isotherm *f_R_*(*R*) as follows:

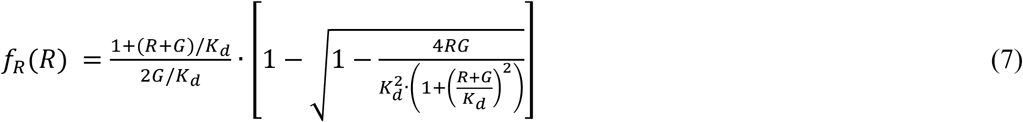

The FRET signal vs *R* can be fit to the expression:

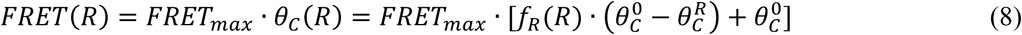

where *θ_C_^0^* is the fraction of C-Gag in the free state, and *θ_C_^R^* is the fraction of C-Gag present in the RNA-bound state, with *f_R_*(*R*) given by eq. (6) or eq. (7).

### Fitting of the Gag dimerization constant

Data for the Gag concentration-dependent FRET change were fit using the following equation:

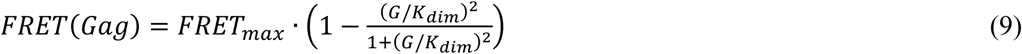

where 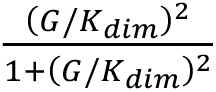 is the fraction of dimerized Gag molecules, and *K_dim_* is the Gag dimerization constant.

### Mass photometry (MP) to probe Gag-RNA complex

MP experiments were performed on a Refeyn Two^MP^ instrument controlled by the Refeyn Aquire^MP^ software. Glass slides were cleaned by washing with alternating Milli-Q water, and 100% isopropanol rinses, for a total of two isopropanol washes, and a final wash with Milli-Q water. The slides were then dried under a stream of nitrogen and gaskets were cleaned following the same strategy but were allowed to air dry for >30 min after removal from the nitrogen stream. Slides were then coated with poly-L-lysine to ensure the surface has a net positive charge. First, a cleaned slide was placed flat on a lint-free surface and 10 μL of a 0.01% poly-lysine solution was placed in the center of coverslip #1. A second cleaned coverslip was placed over the drop perpendicular to the 1st coverslip and they were left to incubate 30-45 sec. The cleaned slides were then separated and dipped into a Falcon tube containing Milli-Q water to remove excess poly-L-lysine. The coated slides were washed twice with Milli-Q water and dried under a stream of nitrogen. To set up for the experiment, a clean gasket was placed in the center of a clean glass slide and then placed in the instrument’s slide holder. First, 19.5 μL of MP reaction buffer (20 mM HEPES pH 7.5, 1 mM βME, 100 mM NaCl, 2 mM Tris-HCl, 1 mM MgCl_2_, and 0.1 ZnCl_2_) was added to gasket well #1 and focus was found. Data for calibration was acquired daily using a mixture of thyroglobulin (monomer and dimer species) and β-Amylase (monomer, dimer, and tetramer species) with known molecular weights. An RNA calibration was also carried out following the same protocol with *in-vitro* transcribed RNAs of varying molecular weight (42.6 kDa, 96.6 kDa, 142.4 kDa, and 254.5 kDa) provided by the lab of Dr. Venkat Gopalan (Ohio State University). The RNAs were run separately following the same procedure. MP reaction buffer was added and focus found as described above, then the RNA was added to the 19.5 μl already present in the gasket well to a final concentration of 5 nM. A 1-min movie was then acquired. The measured ratio-metric contrast values were used to calculate the molecular weight using Refeyn Discover^MP^ software.

Prior to MP measurements, RNA (Psi and TARpA) was refolded at a concentration of 2 µM in 50 mM HEPES pH 7.5 by incubating at 80 °C for 2 min, 60 °C for 2 min, addition of 2.5 mM MgCl_2,_ 37 °C for 30 min and then incubated on ice for >15 min. Gag protein stocks were diluted to 5 µM with Gag storage buffer lacking glycerol (20 mM Tris-HCl pH 7.5, 500 mM NaCl 10 mM βME, and 1 µM ZnCl_2_). RNA-Gag complexes were formed by incubating 500 nM RNA with 500 nM Gag in MP reaction buffer at room temperature for 5 min and then placed on ice. Complexes were diluted in the MP well to 20 nM and a 1-min movie was recorded, following the protocol described above for finding focus. Data were displayed as histograms generated by taking the average number of counts per subpopulation for each RNA species.

### All-atom molecular dynamics (AAMD) simulations

#### System set-up

An atomic model of full-length Gag was constructed using cryo-electron tomography density map-derived atomistic structures of immature MA hexamer of trimers (PDB ID: 7OVQ) (Qu et al. 2021) and CA-Spacer Peptide 1 (SP1) (PDB ID: 5L93, 6BHR) (Dick et al. 2018), and NMR structures of the NC (PDB ID: 1A1T) (De Guzman et al. 1998; Amarasinghe et al. 2000) and p6 (PDB ID: 2C55) domains (Fossen et al. 2005). Myristoylated (Myr) tails, in an exposed conformation, were covalently attached to the N-terminal Gly1 residue using the CHARMM-GUI input generator (Lee et al. 2016). To obtain the initial structure of extended Gag monomers, a membrane-bound dimeric Gag structure with implicit Gag-RNA bonding was initially equilibrated using previously described methods (Banerjee and Voth 2024), and monomers were extracted from the dimeric structure. We used homology modeling tools available in the Rosetta software suite (Watkins et al. 2020) to derive a 98-nt Psi RNA atomistic model using the sequence reported (Webb et al. 2013) together with the atomistic structures (PDB ID: 1A1T, 1F6U) of stem-loop 2 (SL2) and SL3 reported using NMR (De Guzman et al. 1998; Amarasinghe et al. 2000). The RNA-bound monomeric Gag structures were constructed using NC-SL2 and NC-SL3 NMR-structures (De Guzman et al. 1998; Amarasinghe et al. 2000), solvated in 0.15 M KCl and 0.02 M MgCl_2_. The initial box lengths were 19.2 nm × 19.5 nm × 24.0 nm to allow enough water layers to be present between the protein units and the periodic image of the outer leaflet. The total size of each system was ∼ 1 M atoms. These simulations of Psi RNA-bound Gag monomers are denoted as (+)Psi RNA system throughout the manuscript. We prepared another system of Gag monomer alone solvated in water and under the same ionic strength conditions, which is denoted as (-)RNA system in the manuscript. The simulation box length and number of atoms were similar to the (+)Psi RNA system.

#### Simulation details

We used the CHARMM36m force field (Huang et al. 2017) for protein and RNA, and CHARMM TIP3P for water (MacKerell Jr et al. 1998). MD simulations were performed with periodic boundary conditions in all directions using GROMACS 2020.4 MD software (Abraham et al. 2015). Energy minimization was performed using the steepest descent algorithm until the maximum force was less than 1,000 kJ mol^−1^nm^−1^. Equilibration with harmonic restraints (using a 1,000 kJ mol^−1^nm^−2^ spring constant) was carried out on heavy atoms of protein and RNA for 1 ns in the constant NVT ensemble (in which the number of particles (N), volume (V), and temperature (T) remains constant, i.e., canonical ensemble) with a time step of 1 fs, followed by a 15 ns constant NVT MD run with a time step of 2 fs and then a 15 ns MD run in the constant NPT ensemble (in which the number of particles (N), pressure (P), and temperature (T) are held constant, i.e., isothermal–isobaric ensemble) with a time step of 2 fs. The previous harmonic positional restraints were removed, and 1 μs trajectories were generated for 6 replicas for (+)Psi RNA-Gag monomers system and 3 replicas of (-)Psi RNA-Gag monomers system. Throughout this procedure, the temperature was kept constant at 310.15 K using the Nosé-Hoover thermostat with a 1.0 ps coupling constant (Nosé 1984; Hoover 1985), and the pressure was set at 1 bar and controlled using the Parinello-Rahman barostat. The compressibility factor was set at 4.5×10^-5^ bar^-1^ with a coupling time constant of 5.0 ps (Parrinello and Rahman 1981). Van der Waals interactions were computed using a force-switching function between 1.0 and 1.2 nm, while long-range electrostatics were evaluated using Particle Mesh Ewald (PME) (Darden et al. 1993) with a cutoff of 1.2 nm; hydrogen bonds were constrained using the LINCS algorithm (Hess et al. 1997).

#### Analysis of AAMD simulations

Protein conformational changes were visualized using VMD 1.9.3 (Humphrey et al. 1996). Analyses of the trajectories were performed using GROMACS (Abraham et al. 2015), PLUMED (Tribello et al. 2014), MDAnalysis (Gowers et al. 2016), and VMD Tcl scripts (Humphrey et al. 1996).

## Acknowledgments

We thank Dr. Michael Poirier (Ohio State University, OSU) for helpful discussions regarding the FRET measurements and use of his lab’s fluorometer. We also thank Dr. Kevin Jamison (OSU) for help with the FRET calculations. This research was supported by the National Institute of Allergy and Infectious Disease (NIAID) of the National Institutes of Health (NIH) under award numbers R01 AI153216 (KM-F) and U54 AI170855 (VHW, GAV and KM-F) and T32 GM144293 (to KG). Computational resources were provided by the University of Chicago Research Computing Center, the Beagle-3 computer (NIH award 1S10OD028655), and Frontera (at the Texas Advanced Computing Center, TACC) funded by NSF grant OAC-1818253). MP research was supported by NIH RM1 GM149374 (to VHW).

