## Supplementary Files for "Psi RNA-specific Binding Promotes HIV-1 Gag Conformational Change Critical for Immature Viral Particle Assembly"

^1^Department of Chemistry and Biochemistry, Center for RNA Biology, and ^2^Center for Retrovirus Research, Ohio State University, Columbus, OH ; ^3^Department of Chemistry, Chicago Center for Theoretical Chemistry, Institute for Biophysical Dynamics, and James Franck Institute, The University of Chicago, Chicago, IL 60637; ^4^Resource for Native Mass Spectrometry-Guided Structural Biology, Ohio State University, Columbus, OH, and ^5^School of Chemistry & Biochemistry, Georgia Institute of Technology, Atlanta, GA.

^ equal contributions

Yehong Qiu:

Puja Banerjee:

Kaylee Grabarkewitz:

Vicki Wysocki:

Ioulia Rouzina:

Gregory A. Voth:

**Running Title:** Specific RNA Binding Promotes HIV-1 Gag Extension

**Key Words:** HIV-1 assembly, Gag, Psi RNA, nucleocapsid protein (NC), Förster resonance energy transfer (FRET), molecular dynamics simulations, inositol hexakisphosphate (IP6)

**Supplementary Figure S1**

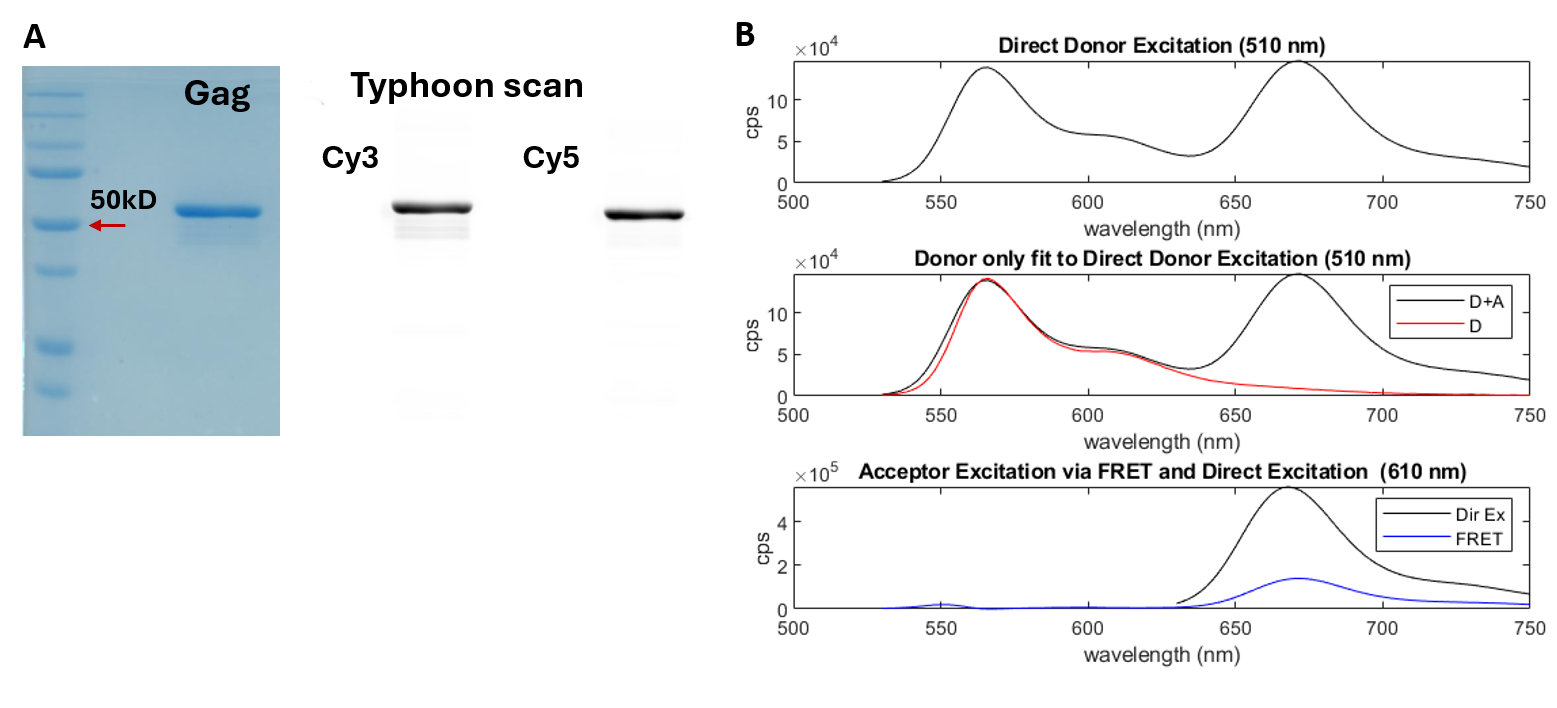

**Figure S1. A) 10% SDS-PAGE gel to assess purity of dual-labeled WT Gag**. Left: Coomassie blue-stained gel; first lane shows molecular weight markers; Right: Fluorescence signal from the same gel. **B) FRET measurement**: Top, measured 510 nm excitation spectrum (530-750 nm); Middle, Cy3-only reference excitation spectrum fitted to the top spectrum to calculate acceptor signal excited by FRET; Bottom, measured 610 nm excitation spectrum (630-750 nm) compared with FRET excitation. The Y-axis is counts-per-second (cps) of photon. The ratio of FRET excitation *versus* direct excitation (bottom) was used to calculate the FRET efficiency using the *(ratio)*_A_ method as described (Clegg 1992).

**Supplementary Figure S2**

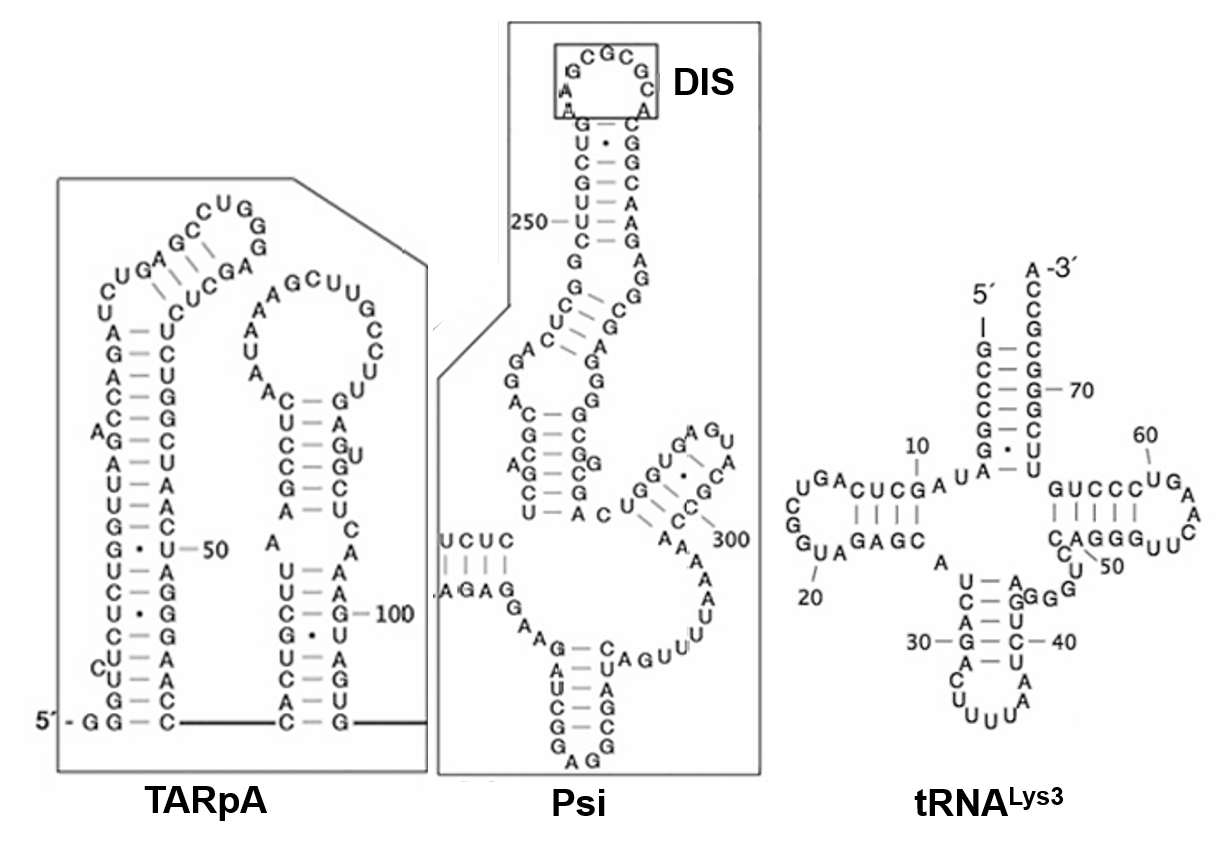

**Figure S2. Secondary structures of RNAs used in this work**.

**Supplementary Figure S3**

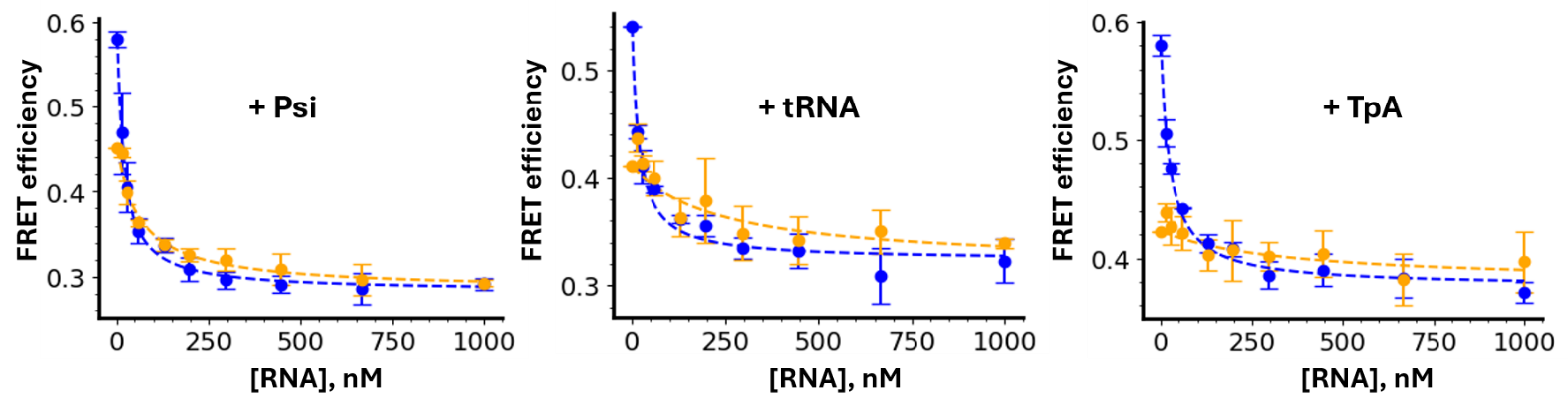

**Figure S3. Overlay of the dual-labeled Gag FRET efficiency *versus* RNA concentration plots for each experimental set obtained with (orange curves) or without (blue curves) 0.3 mM IP6.** Data are the same as in Figures 3 and 5 of the main text.

**Supplementary Figure S4**

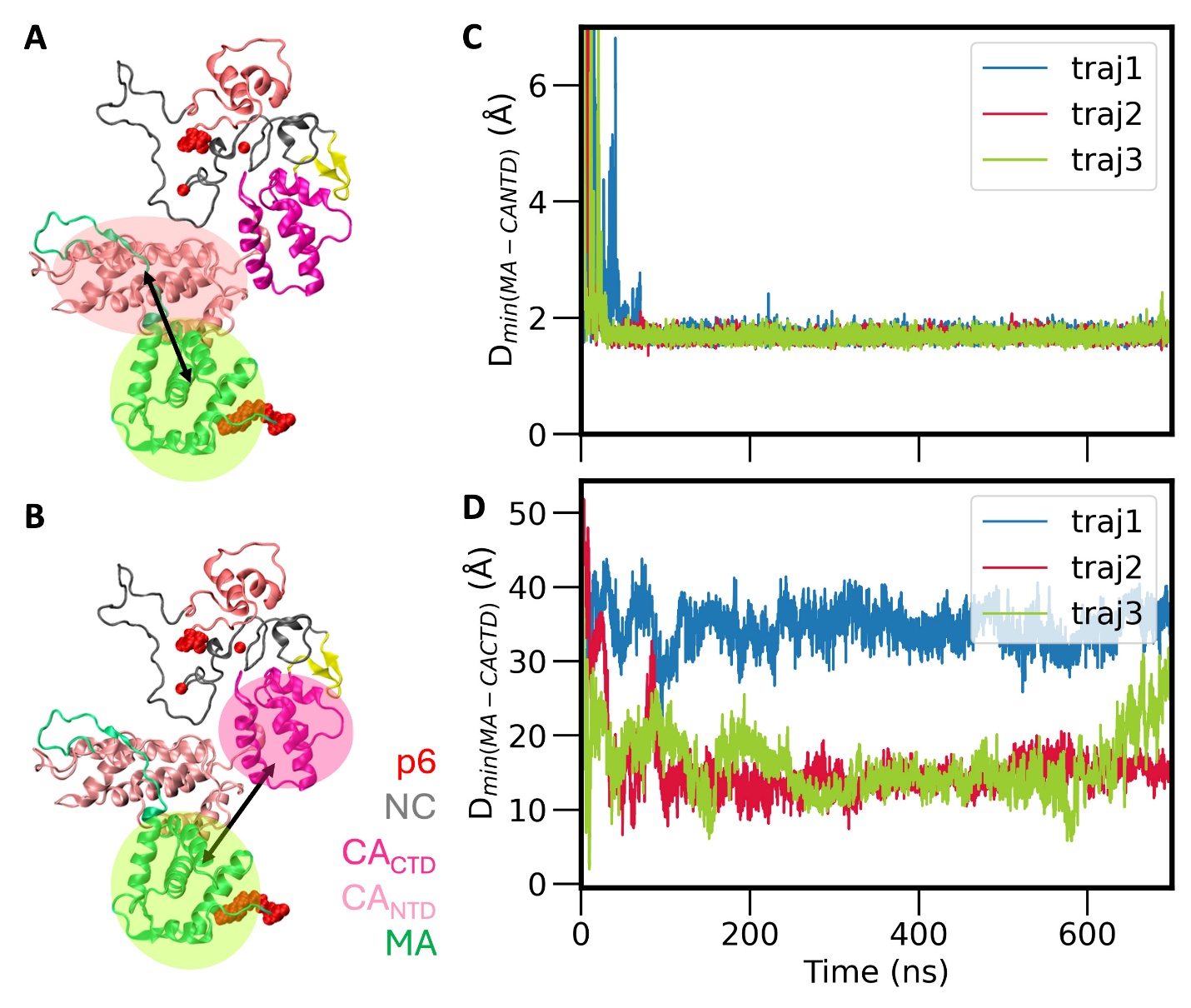

**Figure S4: MA has a higher binding affinity for CA_NTD_ relative to CA_CTD_ in the C-Gag conformation.** (A-B) Representative C-Gag conformations in the (-) RNA system; arrows indicate the distance being measured. (C-D) Time series measuring the minimum distance of the MA domain from the CA_NTD_ (C) and CA_CTD_ (D) domains. MA attains stable binding with the NTD domain in all AAMD trajectories (panel C). MA remains close to the CTD domain in two out of three trajectories (panel D).

**Supplementary Figure S5**

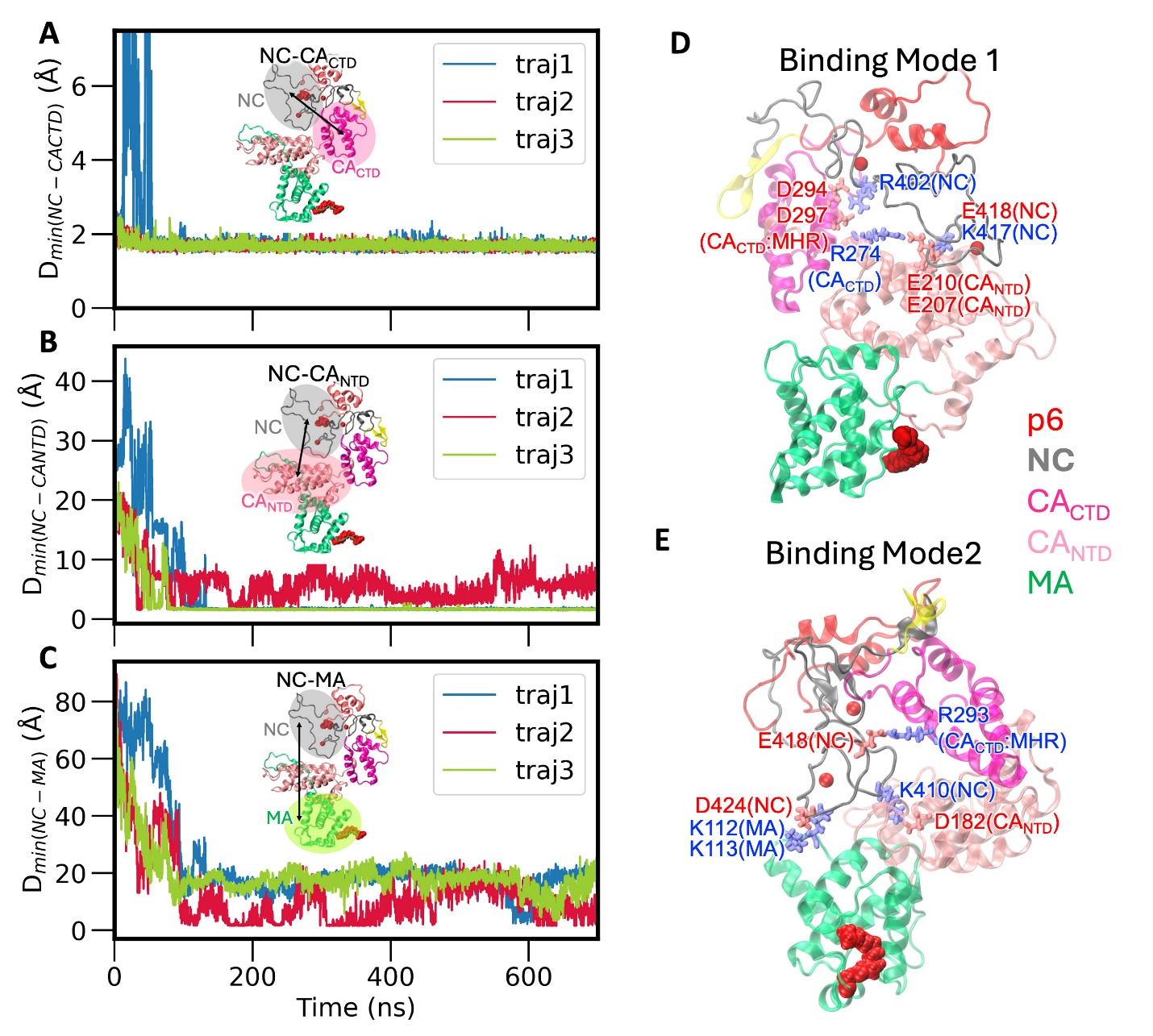

**Figure S5: Nucleocapsid domain (NC) binding to other domains of C-Gag in the absence of RNA.** Time series data of the minimum distance between (A) NC and CA_CTD_, (B) NC and CA_NTD_, and (C) NC and MA. (D-E) Residue-level interactions of NC with CA_NTD_, CA_CTD_, and MA domains in two different binding modes are driven mainly by electrostatic interactions with acidic residues in red and basic residues in blue. In binding mode 1 (panel D), NC does not directly bind to the MA domain. In binding mode 2 (panel E), MA basic residues bind NC acidic residues, as captured by the minimum distance trajectory2 in panel C.

**Table S1** List of primers used in mutagenesis for Gag variants.

| Mutation | Primer | Sequence |
| --- | --- | --- |
| Insert 2 glycine | Forward (tail and short) | 5′-ATG**GGTGGT**GGTGCGAGAGC  GTCGGTATTAAG-3′ |
|  | Reverse (tail and short) | 5′-ACC**ACCACC**CATGGTCTGTTTCC  TGTGTGAAATTGT-3′ |
| Insert 495KCK | Forward (tail and short) | 5′-TT**AAATGCAAA**CCCTCGTCAC  AAGAAAACCTG-3′ |
|  | Reverse (tail and short) | 5′-**TTTGCATTT**AAAGAGTGATCTGA  GGGAAGCTAAAG-3′ |
| Insert W316A, M317A | Forward (tail and short) | 5′-AAT**GCGGCG**ACAGAAACCTTG  TTGGTCCAAAATGCG-3′ |
|  | Reverse (tail and short) | 5′-TGT**CGCCGC**ATTTTTTACCTCT  TGTGAAGCTTGCTCG-3′ |

The tail sequences are underlined with mutations in bold font.

**Table S2** List of RNA sequences used in this study.

| RNA | Length (nt) | sequence |
| --- | --- | --- |
| TARpA | 105 | 5′-GGGUCUCUCUGGUUAGACCAGAUCUGAGCCUGG  GAGCUCUCUGGCUAACUAGGGAACCCACUGCUUAA GCCUCAAUAAAGCUUGCCUUGAGUGCUCAAAGUAGUG-3′ |
| Psi | 107 | 5′-UCUCUCGACGCAGGACUCGGCUUGCUGAAGCGCGCACGG CAAGAGGCGAGGGGCGGCGACUGGUGAGUACGCCAAAAAUUUUGACUAGCGGAGGCUAGAAGGAGAGA-3′ |
| tRNA^Lys3^ | 76 | 5′-GCCCGGATAGCTCAGTCGGTAGAGCATCAGACTTTTAA TCTGAGGGTCCAGGGTTCAAGTCCCTGTTCGGGCGCCA-3′ |
